# Birth by Cesarean Section Remodels Neonatal Brain Barriers and Associated Immune and Metabolic Pathways in Mice

**DOI:** 10.64898/2026.01.06.696394

**Authors:** Valentine Turpin, Jennifer Morael, Alexandre J. C. Cergneux, Hugo Blair, Elisa Cintado, Atreye Majumdar, Carmen Tessier, Lorena Morales, Emily G. Knox, Anna Ratsika, Katherine E. Guzzetta, Ana Paula Ventura-Silva, Jennifer Shearer, Neil J. Loftus, Emily G. Armitage, Alan Barnes, Gerard Clarke, Jonathan R. Swann, John F. Cryan, María R. Aburto

## Abstract

Delivery mode shapes early-life physiology through multiple pathways, including stress and hormonal signalling, neonatal oxygenation, and exposure to maternal microbes. Cesarean section (CS) bypasses several of these processes and has been associated with altered immune maturation and neurodevelopmental outcomes in humans and animal models. Given that CS is known to disrupt gut barrier integrity and that gut and brain barriers operate as a coordinated network, we investigated whether CS affects early postnatal brain barriers.

Using a mouse model, we compared CS-delivered pups to vaginally born (VB) controls at postnatal days 8-9. Brain barrier permeability and structure were assessed using tracer extravasation and vascular and tight junction analyses, choroid plexus immune populations were profiled, and metabolomics was performed in cortex and plasma.

The blood-brain barrier in the medial prefrontal cortex was mostly preserved. In contrast, the blood-cerebrospinal fluid barrier (BCSFB) exhibited structural alterations in vascular and epithelial tight junction compartments. These changes were accompanied by immune remodelling, including expansion of antigen-presenting border-associated macrophages and reduced parenchymal microglia density. CS delivery increased brain tracer accumulation, consistent with altered barrier function during this developmental window. Metabolomic analyses revealed lower levels of key brain metabolites, including N-acetylaspartate, a marker of neuronal metabolic state.

Together, these findings demonstrate that CS delivery induces early alterations in brain barrier function, BCSFB structure, the immune landscape, and metabolic profiles in mice. This identifies delivery mode as a key perinatal variable shaping neonatal brain physiology and establishes a framework for dissecting how birth context may influence long-term neurodevelopmental trajectories.

**HIGHLIGHTS:**

- Cesarean section (CS) reveals the neonatal BCSFB as a vulnerable interface shaped by birth mode
- CS increases brain tracer accumulation despite largely preserved BBB structural markers at the mPFC
- CS induces region-specific vascular, epithelial, and immune remodeling in the choroid plexus
- Cesarean birth increases the proportion of antigen-presenting choroid plexus macrophages
- CS shifts mPFC metabolic profiles, including reduced N-acetylaspartate and malate

## 1. INTRODUCTION

Cesarean section (CS) is a lifesaving surgical intervention whose global use has increased substantially over the last decades^1,2^. While CS remains essential in many clinical contexts, a growing body of epidemiological and pre-clinical evidence has associated delivery by CS with immune, metabolic, and neurodevelopmental outcomes in offspring^3–9^.

Compared with vaginal birth (VB), cesarean delivery, particularly pre-labor (elective) CS, bypasses labor-associated activation of fetal and neonatal glucocorticoid and catecholaminergic pathways, modifies neonatal oxygenation at birth, and prevents direct exposure to maternal vaginal and intestinal microbiota^8,10–12^. These factors play critical roles in postnatal immune education, metabolic adaptation, and tissue maturation of different organ systems. In particular, the absence of maternal microbial transfer has been consistently associated with delayed immune maturation, altered microbial succession, and increased susceptibility to inflammatory and metabolic disorders later in life^4,13–15^. Importantly, many of these birth-associated hormonal, microbial, and immune signals converge on epithelial and endothelial barrier interfaces, suggesting that these structures could represent early targets of CS-associated perturbations. The early postnatal period represents a critical window of vulnerability and plasticity for multiple organ systems, including the immune system and the developing brain. In mice, the end of the first postnatal week is characterized by intense synaptic pruning, neuronal network consolidation, and elevated microglial phagocytic activity, such that disruption of these tightly coordinated processes can lead to altered brain developmental trajectories and long-term behavioral deficits^16–18^. Consistent with this developmental sensitivity, cesarean delivery has been associated with early-life behavioral and physiological alterations in mice, including deficits in early-life and adult social communication, and changes in gut microbiota composition^7^. More recently, CS has also been linked to transient gut barrier dysfunction at postnatal day (P) 7, but not at later stages^19^, indicating that barrier systems are particularly sensitive to peripheral signals, including those associated with delivery mode. This observation is especially relevant given the close cellular and molecular similarities between gut and brain barriers^20^, suggesting that they may be modulated by similar peripheral signals, including those derived from the gut microbiota.

Brain barriers are key players in preserving brain homeostasis. They restrict harmful signals while regulating molecular exchange between the periphery and the brain, primarily at two key interfaces: the blood-brain barrier (BBB) and the blood-cerebrospinal fluid barrier (BCSFB). The BBB relies on the neurovascular unit (NVU), a dynamic cellular assembly of endothelial cells, connected by both tight and adherens junctions, as well as pericytes, astrocytes, and microglia, that collectively regulate BBB function and permeability^20,21^. Passive and active transport of molecules is limited by specific transporters, a limited rate of transcytosis, and, more importantly, tight junctions such as Claudin-5, Occludin, and Zonula occludens-1. Importantly, NVU assembly and BBB maturation undergo extensive postnatal remodeling, resulting in heightened plasticity and increased vulnerability to perinatal challenges, with potentially lasting consequences for brain homeostasis and neurodevelopmental outcomes^22,23^.

The ChPs, located within all brain ventricles [lateral (LV), third (3V), and fourth (4V)], represent a second major interface controlling the exchanges between the blood and the CSF through the BCSFB. This barrier plays a central role in regulating CSF composition, thereby directly influencing neurodevelopmental processes and overall brain function^24–26^. Permeability of the BCSFB is primarily governed by epithelial cells, which are sealed by tight junction complexes analogous to those in the BBB, restricting the passage of small- and medium-sized molecules^27^. Additionally, the ChP endothelium is composed of fenestrated vessels, which allow the passage of larger molecules. Notably, this vascular interface can dynamically restrict permeability in inflammatory contexts^28^. In addition to its barrier function, the ChP constitutes a specialized neuroimmune niche populated by diverse immune cell types, including lymphocytes and macrophages^29^. Among them, ChP-resident macrophages, also called border-associated macrophages (BAM), protect the brain from peripheral insults as early as embryonic day 9^30,31^.

Increasing evidence indicates that brain barriers actively participate in shaping neurodevelopmental trajectories under both physiological and pathological conditions^20,23,32,33^. Notably, the BCSFB has emerged as a key immunological gateway during development, capable of rapidly responding to peripheral inflammatory signals and modulating cerebrospinal fluid composition^26^. Despite this, most studies examining perinatal insults have focused on the BBB or on later developmental stages, leaving the early postnatal BCSFB largely unexplored as a potential site of vulnerability.

Both brain barriers become functional from mid-embryogenesis, underscoring their role in maintaining brain homeostasis during periods of heightened developmental vulnerability^23,34,35^. These features make them plausible targets for perinatal peripheral influences capable of shaping neurodevelopmental trajectories.

Based on this rationale, we hypothesized that cesarean delivery disrupts the maturation and function of the BBB and BCSFB during the early postnatal window and evaluated this using complementary in vivo permeability assays and high-resolution confocal imaging.

## 2 MATERIALS AND METHODS

### 2.1 Animal husbandry

All procedures were approved by the Animal Experimentation Ethics Committee (AEEC) at University College Cork and the Health Products Regulatory Authority (HPRA). Experiments were conducted under project authorization number AE19130/P169 in accordance with the recommendations of the Directive 2010/63/EU.

8-week-old NIH Swiss were fed ad libitum and housed individually in a 12-hour day/night light cycle, temperature 22°C (+/-2) and humidity 55% (+/-10). After pairing, the detection of a vaginal plug was recorded as gestational day 0.5 (G0.5) and females were separated and housed individually. Once the pregnancy was confirmed, females were attributed to the cesarean section (CS) or vaginal birth (VB) groups.

### 2.2 Cesarean section procedure

The CS procedure was performed as previously described in Morais et al., 2020^7^. Briefly, at day 19.5, pregnant females were euthanized by cervical dislocation, and pups were extracted and massaged until spontaneous breathing was observed. They were then given to a foster dam who had vaginally delivered within the previous 24 hours. Detailed procedures are provided in the SI Appendix.

### 2.3 Flow cytometry of the choroid plexus

P8 mice were euthanized by anesthetic overdose (pentobarbital 60-90mg/kg) and perfused intracardially with Phosphate Buffered Saline (PBS). Lateral ventricle (LV) and fourth ventricle (4V) choroid plexuses were dissected under stereomicroscope (SMT1, Tritech Research) and placed in PBS with 10% heat-inactivated Fetal Bovine Serum (FBS) (F9665, Bio-Sciences). Each n=1 sample consisted of choroid plexuses from three sex-matched littermates. Tissues were processed and gated as described in the SI Appendix. Flow cytometry outputs included percentage of parent populations, and median fluorescence intensity (MFI) for activation of cell surface markers.

### 2.4 In vivo procedures, sample collection and tissue processing

#### 2.4.1 In vivo brain barrier permeability assessment

At P8, 30µL of Alexa Fluor™ 555 cadaverine 1kDa (2 mg/ml) (Thermo Fischer) was injected intraperitoneally and allowed to circulate for 1 hour. Mice were then euthanized by anesthetic overdose and perfused intracardially with 4% paraformaldehyde (PFA) in PBS. After 1 hour of post-fixation at 4 °C, whole brains mounted in PBS were imaged on the ZEISS SteREO Discovery.V20 with MikroCamII 9MP 4K 1’’ Microscope Camera and associated software (Bresser, GmbH) at 7.5x magnification. Brains were post-fixed at 4 °C for an additional 5 hours after imaging and stored in anti-freeze solution at -20 °C until tissue processing.

Cryoprotection was performed as previously described by Cui et al., 2020^30^. Brains were sectioned into 40-µm sections using a cryostat (Leica CM1900) and mounted onto SuperFrost Plus slides (J1800AMNZ, Epredia). Immunofluorescence protocols, including antibody references, confocal imaging, and subsequent analysis protocols are described in the text of the SI Appendix and Table S1. For each animal (n=1), mean fluorescence intensity corresponds to the mean grey value averaged from 1–6 images (2–3 ROIs per image) for the choroid plexus and from 4–6 images for medial prefrontal cortex sections.

#### 2.4.2 Tight junction and vasculature analysis

P8 mice were euthanized by anesthetic overdose as previously described and perfused intracardially with PBS and then with 10% trichloroacetic acid (TCA). Brains were post-fixed at 4 °C for 6 hours and stored in anti-freeze solution at -20 °C until tissue processing. Brains were either sectioned coronally at 80μm using a vibratome (Leica, VT1000 S) or 40-µm using a cryostat (Leica CM1900). Immunofluorescence protocols, including antibody references, confocal imaging, and subsequent analysis protocols are described in the text of the SI Appendix and Table S1. For each animal (n=1), the results of 6 brain sections were averaged.

### 2.5 Plasma and cortex metabolomics

Two different sample types from P9 NIH Swiss mice were analyzed, each by a different paradigm. Cerebral cortex (n=13 VB, 16 CS) tissue was collected and analyzed by high-resolution liquid chromatography-mass spectrometry (LC-MS/MS) and targeted analysis was performed for 46 metabolites previously found to be related to gut-brain axis signaling (Swann 2020). Plasma (n=13 VB, 16 CS) was collected and analyzed by 1H NMR spectroscopy. NMR spectral profiles were integrated to calculate the relative abundance of 33 plasma metabolites. Metabolite abundance data from both sample types were analyzed via the same common pipeline. Where necessary, missing values were imputed via the k-Nearest neighbor method, and data was normalized by Total Sum Scaling. Principal component analysis was performed followed by PERMANOVA to determine if the variance in the data could be explained by birth mode. To discover individual metabolites that were altered following CS delivery compared to natural birth, Mann-Whitney tests were carried out on each metabolite, with Benjamini-Hochberg (BH) FDR correction applied.

### 2.6 Statistics

Investigators were blinded to the experimental groups during data acquisition and analyses. Results are expressed as mean ± SEM and each data point represent a single animal. Sex was initially included as a biological variable in the statistical models. No significant sex × delivery mode interaction was detected for the assessed outcomes; therefore, data from males and females were pooled for subsequent analyses. Statistical analyses were performed and plotted using GraphPad v10 (San Francisco, USA). For normal samples with equal variances, Student’s t-tests were used to compare differences between the CS and VB groups. For analysis of the percentages of CD68^+^ immune cells within Iba1^+^ cells, two-way ANOVAs followed by an Uncorrected Fisher’s LSD posthoc test were performed. For data sets with normal distribution but unequal variances, we performed unpaired t-tests with Welch’s correction. For non-normal samples, the Mann-Whitney non-parametric test was carried out. Statistical outliers were excluded using Grubbs’s test, and p-values inferior to 0.05 were considered statistically significant.

## 3 RESULTS

### 3.1 Cesarean delivery minimally alters BBB structural readouts but reduces microglial density in mPFC

We first compared the BBB structure in the medial prefrontal cortex (mPFC) of P8 pups delivered by CS to VB pups (Fig.1, A). Since tight junctions are highly dynamic and crucial in BBB integrity, we quantified the expression of the tight-junction genes Claudin 5 (Cldn5), Occludin (Ocln), and Tight Junction Protein 1/Zonula Occludens 1 (Tjp1/ZO-1). ZO-1 expression tended to be lower in CS-delivered pups (p=0.052; Fig. 1, B), whereas Cldn5 and Ocln expression remained unchanged between groups (Fig. 1, B). This trend suggests a potential modulation of tight junction dynamics in CS-delivered pups, warranting further investigation at the protein and functional levels. Therefore, we measured ZO-1 vascular coverage and intensity but observed no difference across delivery modes (Fig. 1, C-D). In addition, ultrastructure assessment of endothelial ZO-1, inspired by recent analytical approaches developed for adherens junctions^36^, showed no differences in the distribution of the five described morphological patterns (Tremble, Twist, Discontinuity, Amalgam, and Mess) between delivery groups (Fig. 1, F-G; SI Appendix, Fig. S1, A-B). Overall, these findings indicate that mode of delivery induces, at most, subtle transcriptional modulation of tight junction components without detectable changes at the protein or structural level.

**Figure 1.**
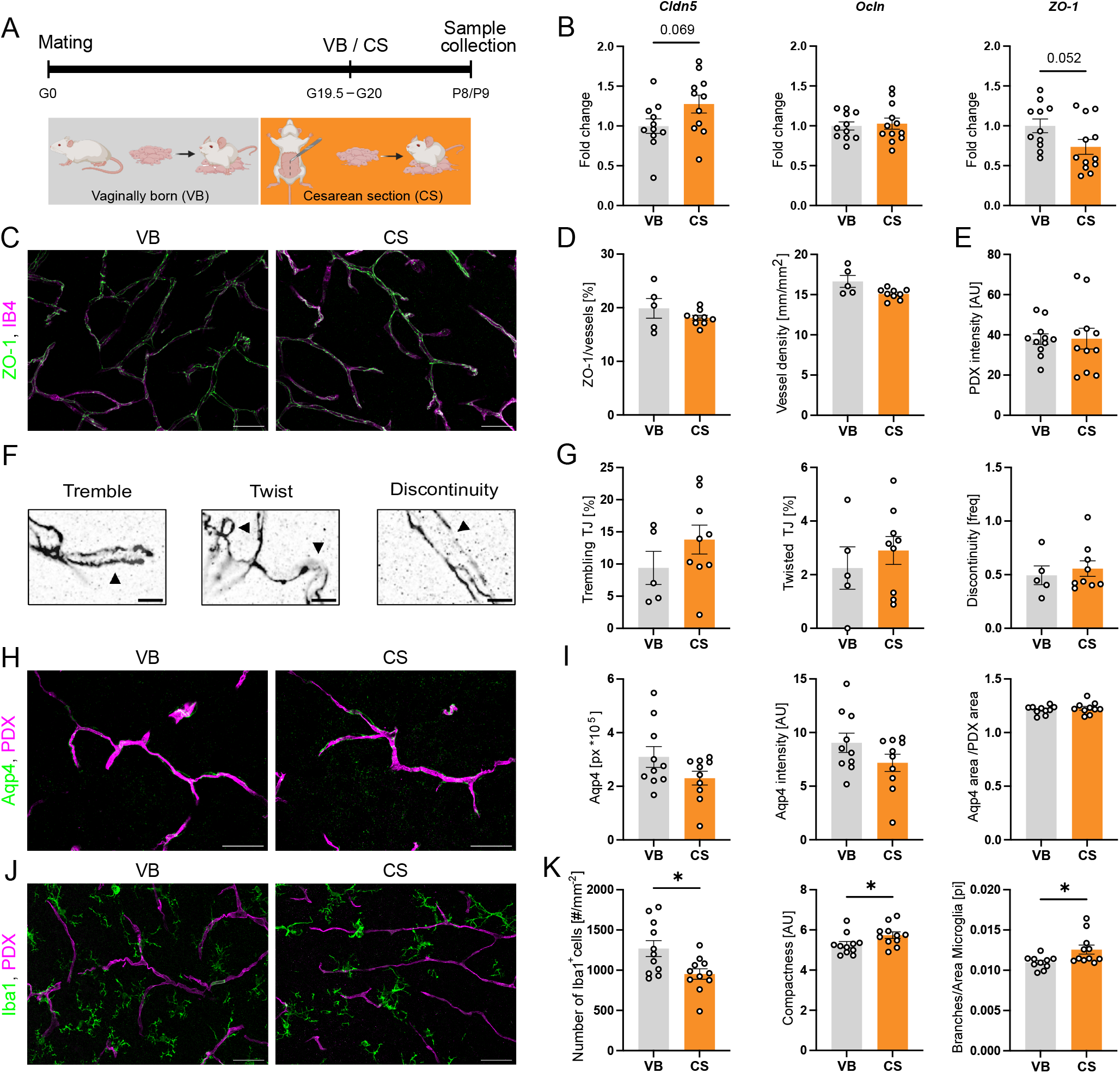
CS minimally alters BBB structural readouts but reduces microglial density in mPFC (A) Schematic overview of the experimental design. (B) Relative mRNA expression of tight-junction genes Occludin (Ocln), Claudin-5 (Cldn5), and Zonula occludens-1 (ZO-1) in the medial prefrontal cortex (mPFC), quantified by qPCR. (mean ± SEM; t-test, n=11-12). (C) Representative images of tight junction (ZO-1) and vessels (IB4) observed in VB and CS mice at P8 (80µm vibratome section, scale bar = 40μm). (D) Fluorescence quantification of ZO-1 over IB4 and vessel density mm/mm2. (mean ± SEM; Welch’s t-test, n=5-9). (E) Fluorescence quantification of the vascular marker PDX (mean ± SEM; Welch’s t-test, n=11). (F) Representative confocal images (Scale bar = 5μm) of tight junction ultrastructure, visualized by ZO-1 immunostaining in cortical microvessels. (G) Analysis of tight junction ultrastructure characteristics: percentage of trembled or twisted TJ in the vessels, frequency of discontinuity in the TJ network (mean ± SEM; t-test, n=5-9). (H) Representative images of astrocytic feet (Aqp4) and blood vessels (PDX). (40µm cryosections, scale bar = 40μm). (I) Quantitative analysis of astrocytic end feet parameters: Aqp4 signal (in pixels), Aqp4 fluorescence mean intensity, percentage of colocalization of Aqp4 and vessel marker. (mean ± SEM; t-test, n=10). (J) Representative images of microglia (Iba1) and blood vessels (PDX) in the mPFC. (40µm cryosections, scale bar = 40μm). (K) Quantitative analysis of microglial parameters: microglia density, number of branches/area and compactness (mean ± SEM; t-test, * p<0.05, n=11).

The analysis of the vascular network in the mPFC demonstrated no effect of delivery mode on vessel density or any of the analyzed vascular properties (Fig. 1, C-D, SI Appendix, Fig. S1, C). The intensity of vascular podocalyxin (PDX), a major constituent of the glycocalyx, was not affected either (Fig. 1, E). Similarly, analysis of Plasmalemma Vesicle-Associated Protein (PV1) fluorescence intensity, a marker of endothelial fenestrations, showed no differences between groups (SI Appendix, Fig. S1, D). This indicates that endothelial fenestration markers, vascular glycocalyx, and general vascular morphology are preserved in the mPFC upon CS at P8.

Finally, we examined astrocytes and microglia, which play key roles in the regulation of BBB structure and neuroimmune signaling. Particularly, astrocytic end-feet, marked by the water channel Aquaporin-4 (Aqp4), are key regulators of BBB maintenance. Several parameters including total Aqp4 signal, Aqp4 intensity, and Aqp4/PDX colocalization showed no difference between groups (Fig. 1, H-I). In contrast, we found a significant reduction in the total microglia density in CS animals (p<0.05, Fig. 1, J-K). However, the number of vessel-associated microglia (VAMs), known to be crucial for vascular remodeling and plasticity^37,38^, and their proportion within the Iba1+ cell population and general Iba1 intensity were not affected by mode of delivery (SI Appendix, Fig. S1, E). Notably, morphological analysis revealed a significant increase in microglial compactness index (measure of cell shape irregularity) and in branching density normalized to cell area in CS animals compared to VB controls (p<0.05, Fig. 1, J-K).

Together, these findings indicate that, despite preserved structural organization of astrocytic end-feet, CS is associated with reduced microglial abundance and a shift in microglial morphology characterized by altered branching density.

### 3.2 Mode of delivery modulates BCSFB endothelial and epithelial readouts in a region-specific manner

The gut and blood-brain barriers have received much more attention than the BCSFB in the context of early-life exposures. Thus, we next assessed the integrity of the BCSFB at the ChP in P8 CS-delivered and VB pups. We first analyzed ChP vessel organization and structure in both the LV and 3V ChPs (Fig. 2, A and D). There was a trend towards increased PDX fluorescence (p=0.0584; Fig. 2, B-top) in CS-born pups in the LV ChP but no changes were detected in vascular organization (Fig. 2, C-top; SI Appendix, Fig. S2, A) or PV1 fluorescence intensity (Fig. 2, D and E-top).

**Figure 2.**
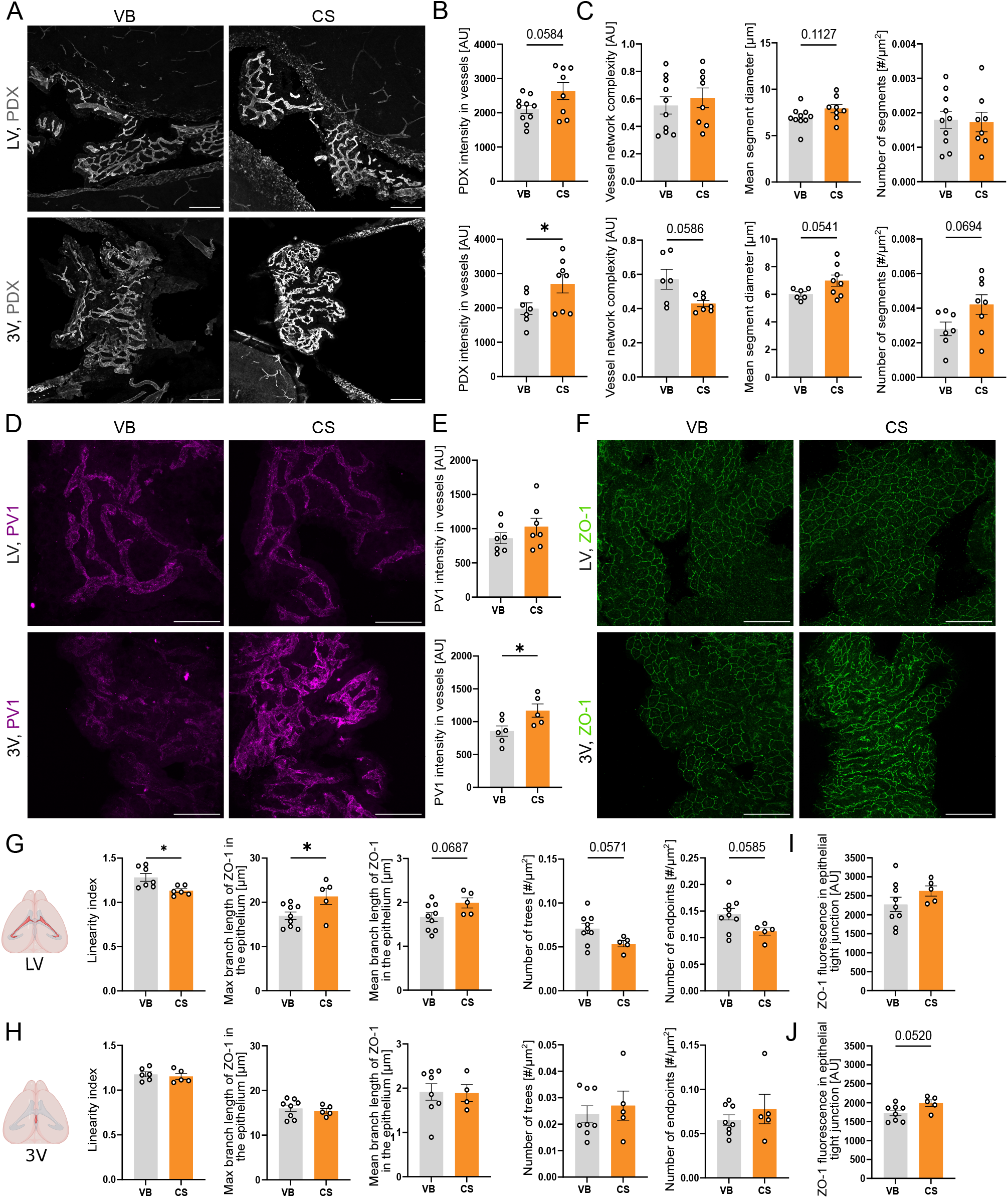
CS affects vascular and epithelial integrity of the BCSFB (A) Choroid plexuses PDX fluorescence observed in VB and CS mice at P8 (40µm cryosections, scale bar = 100μm). (B-E) Fluorescence quantification (B) and vessel network analysis outputs (vessel network complexity, mean segment diameter, and number of segments) (C) in the LV ChP (B-top and C-top) and in the 3V ChP (B-bottom and –C-bottom) (mean ± SEM ; t-test or Welch’s t-test *p<0.05, n=8-10 in the LV ChP, n=6-8 in the 3V ChP). (D and F) PV1 and ZO-1 (F) fluorescence observed in VB and CS mice at P8 (40µm cryosections, scale bar = 50 μm). Fluorescence quantification of PV1 in the LV ChP (E-top) and in the 3V ChP (E-bottom) (mean ± SEM; t-test *p<0.05, n=7 in the LV ChP, n=5-6 in the 3V ChP). (G-J) Fluorescence quantification of ZO-1 (I and J) and the tight junction epithelial network analysis outputs (max branch length, mean branch length, number of trees, number of branches, number of endpoints) (G and H) in the LV ChP (G-I) and in the 3V ChP (H-J). (mean ± SEM; t-test or Mann-Whitney test *p<0.05, n=4-9 in the LV ChP, n=4-8 in the 3V ChP). (A, D and F) Upper panels: LV ChP; bottom panels: 3V ChP; left panels: VB; right panels: CS. PDX: podocalyxin, PV1: plasmalemma-vesicle associated protein 1, ZO-1: Zonula-occuldens-1, VB: Vaginal birth, CS: cesarean section, LV ChP: lateral ventricle choroid plexus, 3V ChP: third ventricle choroid plexus.

In contrast, the strongest changes were observed in the 3V ChP, where fluorescence intensity of both PDX and PV1 was significantly increased in CS-born pups (p<0.05, Fig. 2, A and B-bottom, and Fig.2 D and E-bottom, respectively). These observations revealed that CS delivery modulates structural organization and fenestrated endothelial phenotype of ChP vessels. We then assessed different properties of the vascular network within the 3V ChP. Vessel network analyses showed no significant differences in overall network parameters (SI Appendix, Fig. S2, B). However, mean segment diameter showed a trend towards an increase (p=0.0541), while network complexity tended to be decreased (p=0.0586) in CS-born compared to VB pups (Fig. 2, C-bottom). Unlike the vascular barrier, which is mostly permissive and only closes in specific contexts^28^, the epithelial component of the BCSFB is highly restrictive and only permeable in certain pathological conditions^29,39^. To assess epithelium integrity, we performed a skeleton network analysis after immunostaining for ZO-1 (Fig. 2, F). As previously reported, we observed different modulations of the LV and 3V ChPs. In the LV ChP, CS was associated with a decreased tight junction linearity index (p<0.05, Fig. 2, G) which was not observed in the 3V ChP. This indicates that LV ChP epithelial tight junctions take on a more linear organization compared to VB pups. CS-born pups displayed an increase in maximum branch length of ZO-1 in the epithelium of the LV ChP compared to VB pups (p<0.05, Fig. 2, G), while no changes in the tight junction protein network were observed in the 3V ChP (Fig. 2, H, SI Appendix, Fig. S3, B). No changes were observed for the mean branch length of the tight junction network in the LV ChP (p=0.0687; Fig. 2, G) or in the 3V ChP (Fig. 2, H), nor for other tight junction network properties (SI Appendix, Fig. S3, A-B). However, trends towards reduced tight junction network complexity were observed, including reductions in the number of trees and endpoints in the LV ChP (p=0.0571 and p=0.0585, respectively; Fig 2, G). Analysis of ZO-1 fluorescence intensity showed no significant differences between CS and vaginal delivery in both ChPs, except for a trend towards an increase in the 3V ChP (p=0.0520; Fig. 2, I and J). Taken together, these results demonstrate that cesarean delivery impacts the integrity of the vessel and tight junction networks and differentially modulates the LV and 3V ChPs.

### 3.3 Cesarean delivery affects choroid plexus immune status in a region-specific manner

The ChP, in addition to controlling exchanges between the blood and the CSF, acts as an immune barrier protecting the brain from blood-borne pathogens and inflammatory signals. To assess whether mode of delivery exerts an effect on ChP immune status, we quantified the numbers of both Iba1- and CD68-positive cells which are representative of activated and phagocytic macrophages^30,40^ (Fig. 3, A). In the LV ChP, we observed an increase in the number of Iba1^+^ cells in CS-born pups compared to VB pups (p<0.05; Fig. 3, B) while the number of CD68^+^ cells remained unchanged. In the 3V ChP, there were no changes in either the number of Iba1^+^ or CD68^+^ cells between CS-delivered and VB pups (Fig. 3, C). The percentage of CD68^+^ cells within the Iba1^+^ population was unchanged in the LV ChP but was significantly decreased in the 3V ChP in the CS group compared to the VB group (p<0.05), Fig. 3, C). Iba1^+^ cells located outside of the ChP borders of the ChPs were considered epiplexus macrophages, which are tissue-resident macrophages of the ChP. Their numbers were unchanged in CS-delivered pups compared to VB pups in both LV and 3V ChPs (Fig. 3, B-C).

**Figure 3.**
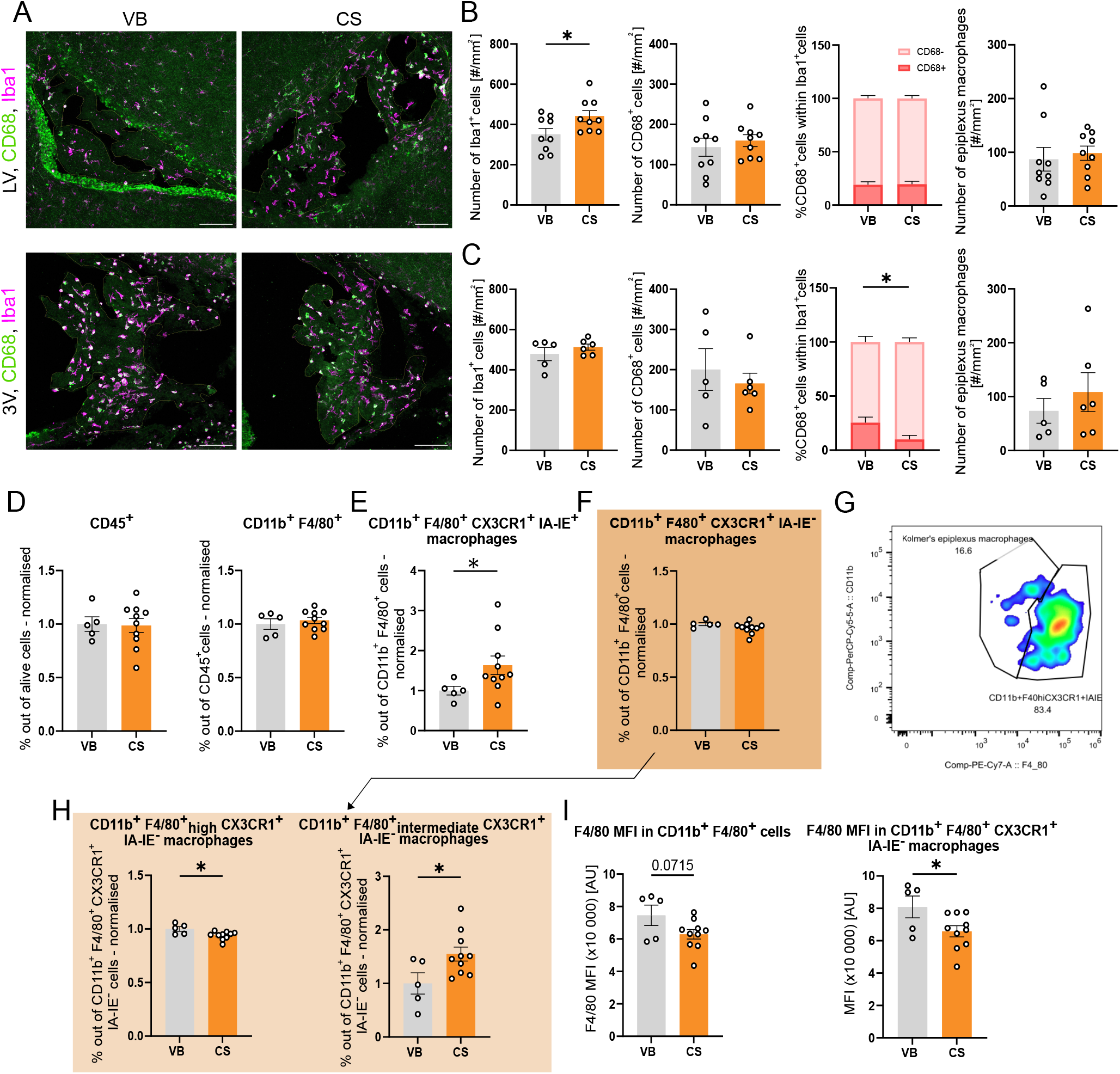
CS impacts ChP immune status through the reorganization of its immune cell populations (A) Choroid plexuses fluorescence observed in VB and CS mice at P8 after CD68 and Iba1 staining (40µm cryosections, scale bar = 100 μm). Upper panels: LV ChP; bottom panels: 3V ChP; left panels: VB; right panels: CS. (B-C) Immune cell counting (Iba1+, CD68+, and epiplexus macrophages) and percentages of Iba1+ CD68+ and Iba1+ CD68-cells in the LV ChP (B) and in the 3V ChP (C) (mean ± SEM ; t-test, Mann-Whitney or 2-way ANOVA *p<0.05, n=8-9 in the LV ChP, n=5-6 in the 3V ChP). (D) Percentages of parent population for the CD45+ and CD11b+ F4/80+ (E-F) Percentage of parent for the CD11b+ F4/80+ CX3CR1+ IA-IE+ (E) and CD11b+ F4/80+ CX3CR1+ IA-IE-(F) macrophages (mean ± SEM; t-test or Welch’s t-test *p<0.05, n=5-10). (G) Gating strategy used to differentiate Kolmer’s epiplexus macrophages from highly expressing F4/80 BAM (x-axis: F4/80; y-axis: CD11b). (H) Percentage of parent for the CD11b+ F4/80+high CX3CR1+ IA-IE-macrophages and CD11b+ F4/80+ intermediate CX3CR1+ IA-IE-macrophages (Kolmer’s epiplexus macrophages). Daughter populations of (F). (mean ± SEM; t-test *p<0.05, n=5-10) (I) F4/80 MFI of the CD11b+ F4/80+ and CD11b+ F4/80+ CX3CR1+ IA-IE-macrophages (mean ± SEM; t-test *p<0.05, n=5-10). Iba1: Allograft Inflammatory Factor 1, CD11b: Cluster of Differentiation 11b, CD45: Cluster of Differentiation 45, CD68: Cluster of Differentiation 68, F4/80: EGF-like module-containing mucin-like hormone receptor-like 1, IA-IE: I-A and I-E α/β heterodimeric major histocompatibility complex II molecules, CX3CR1: CX3C motif chemokine receptor 1, BAM: Border-Associated Macrophages, MFI: Median Fluorescence Intensity, VB: Vaginal birth, CS: Cesarean section, LV ChP: Lateral ventricle choroid plexus, 3V ChP: Third ventricle choroid plexus.

We next performed flow cytometry of the ChPs to complement immunofluorescence results (Gating in SI Appendix, Fig. S4). We did not observe any differences regarding the percentage of CD45^+^ cells, indicating that the number of immune cells was unchanged in CS-born pups compared to VB pups (Fig. 3, D). We then assessed the IA-IE-positive and -negative macrophage populations, which differentiate IA-IE^+^ antigen-presenting cells (MHCII^+^) from IA-IE^-^ non-antigen presenting cells (MHCII^-^). As shown in Figure 3, E, CS led to an elevated percentage of IA-IE^+^ macrophages (CD45^+^ CD11b^+^ F4/80^+^ CX3CR1^+^ IA-IE^+^; p<0.05), whereas the IA-IE-population was unchanged (Fig 3. F). Finally, two populations of border-associated macrophages (BAMs) were analyzed as a function of their expression of F4/80, a marker of murine immune cell maturation. As previously described by Dominguez-Belloso and colleagues^41^, Kolmer’s epiplexus macrophages have a more intermediate expression of F4/80 (CD45^+^ CD11b^+^ F4/80^+intermediate^ CX3CR1^+^ IA-IE^-^; Fig. 3, G). This population was significantly increased in the choroid plexus of CS-delivered pups compared to VB pups (p<0.05; Fig. 3, H). In addition, F4/80^high^ BAMs (CD45^+^ CD11b^+^ F4/80^+high^ CX3CR1^+^ IA-IE^-^) showed a decrease in their number in CS-born pups compared to VB pups (p<0.05; Fig. 3, H), which hints towards a switch in both CD45^+^ CD11b^+^ F4/80^+^ CX3CR1^+^ IA-IE^-^ populations. Interestingly, the median fluorescence intensity (MFI) of the F4/80 cell surface protein showed a decrease in the CS group in the IA-IE^-^ population compared to the VB group (CD45^+^ CD11b^+^ F4/80^+^ CX3CR1^+^ IA-IE^-^; p<0.05; Fig. 3, I). Together, these data suggest that cesarean delivery induces a complex and ChP-specific reorganization of the different populations of immune cells.

### 3.4 Mode of delivery alters functional brain permeability and cerebral cortical metabolic profiles

To determine whether the subtle CS-associated changes observed at the level of the BBB, or the alterations detected at the BCSFB, resulted in any functional permeability deficits, we assessed barrier integrity at P8 by evaluating the distribution of a 1 kDa fluorescent tracer (Alexa Fluor 555–cadaverine) following intraperitoneal injection. Whole-brain imaging revealed increased tracer extravasation measured in the cortex, hypothalamus, and cerebellum of CS-delivered pups (p<0.0001, Fig. 4, A). To further characterize tracer distribution at candidate brain interfaces, brains were sectioned, and the mPFC and ChP were imaged as representative sites of the BBB and BCSFB, respectively. In the mPFC, cadaverine fluorescence was higher in the parenchyma of CS animals, indicating increased functional tracer access to the parenchyma at this site despite preserved BBB structural markers (p<0.05, SI Appendix, Fig. S5, A-B). In the LV ChP, no differences were detected in tracer abundance within either the stroma or the vasculature (SI Appendix, Fig. S5, C-D). In contrast, in the 3V ChP, cadaverine fluorescence was increased in the vascular compartment of CS animals (p<0.05), but not in the stromal region at this time point (SI Appendix, Fig. S5, E-F).

**Figure 4.**
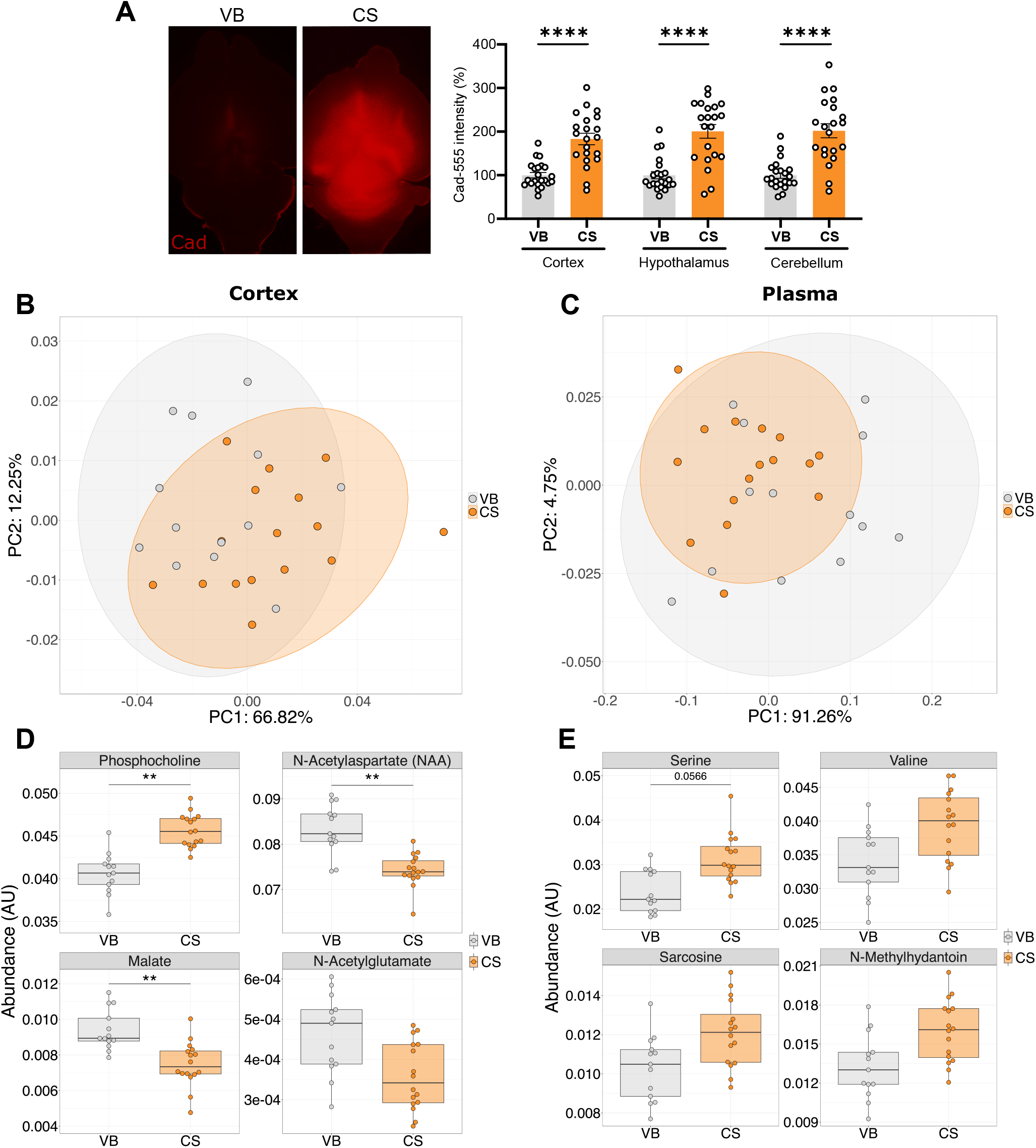
CS alters plasma and cortex metabolomic signatures and is associated with increased functional tracer accumulation in the whole brain (A) Representative dorsal view images of brains and fluorescence intensity percentage of several brain areas (cortex, hypothalamus, cerebellum) of P8 pups vaginally-born (VB) and delivered by cesarean section (CS) after an in vivo Cadaverine Alexa Fluor 555 (1kDa) i.p. injection (mean ± SEM ; Mann-Whitney, ***p<0.001, n=21-22). (B) Principal component analysis of metabolomes from VB and CS P9 cerebral cortices. PERMANOVA analysis indicates delivery mode accounts for 15% of the variance between groups (p<0.05). (C) Principal component analysis of metabolomes from VB and CS P9 plasma. PERMANOVA analysis indicates delivery mode accounts for 14% of the variance between groups (p<0.05). (D) Relative abundance of the top 4 most significantly altered metabolites in cerebral cortex (Mann-Whitney U-Test with BH FDR **p<0.01). (E) Relative abundance of the top 4 most significantly altered metabolites in plasma (Mann-Whitney U-Test with BH FDR).

Finally, metabolomic analyses were performed on cortical tissue and plasma to explore potential metabolic alterations associated with the observed brain barrier phenotypes. Using liquid chromatography-mass spectrometry (LCMS), a targeted panel of 46 metabolites was measured in cerebral cortex samples collected at P9. Similarly, the metabolic profiles of plasma sampled at P9 were measured by 1H nuclear magnetic resonance (NMR) spectroscopy and the relative abundance of 33 metabolites was extracted from the spectra. Through principal component analysis and PERMANOVA, we examined changes in cerebral cortex and blood plasma profiles at P9 as a consequence of delivery mode. In both samples, the profiles were subtly but significantly altered, explaining 14% of the variance in the cortex (p < 0.05; Fig. 4, B) and 15% in plasma (p < 0.05; Fig. 4, C). In the cortex, three metabolites were significantly altered, with phosphocholine increased in CS-delivered animals, while N-acetylaspartate (NAA) and malate were decreased compared to VB animals (Mann-Whitney test with BH FDR; q < 0.05; Fig. 4, D). In the plasma, only serine showed a trend towards being significantly altered by mode of delivery (q = 0.057; Fig. 4, E). Collectively, this data strongly indicates brain barrier function impairments and suggests metabolism alterations in relation to CS delivery.

## 4. DISCUSSION

Early-life environmental exposures are increasingly recognized as critical regulators of brain barrier development. Here, we identify mode of delivery as a previously underappreciated perinatal factor that selectively shapes BBB and BCSFB properties during early postnatal life in mice. While CS induced only subtle transcriptional changes in BBB parameters, the BCSFB emerged as a major site of vulnerability, with structural and immune alterations affecting its endothelial, epithelial, and immune compartments. These changes were accompanied by an increased brain tracer signal, with specific region-dependent leakage, and metabolomic signatures in the brain and plasma associated with delivery mode. These results identify CS as a perinatal susceptibility model in which brain barriers are at least transiently affected during a critical developmental window.

While we cannot completely rule out BBB disruption, molecular, ultrastructural, and cellular analyses consistently indicated preserved vascular architecture, intact tight junctions, and normal astrocytic end-foot coverage. Microglial density was reduced in the mPFC, without evidence of classical amoeboid activation or changes in VAMs. However, microglia exhibited increased branching density and higher compactness, indicating a shift in morphology. As classical inflammatory activation is associated with process retraction and reduced ramification, this profile is instead consistent with a primed or reactive state. Such intermediate phenotypes, characterized by increased process complexity, have been described in response to mild perturbations. This contrasts with some models of maternal immune activation (MIA) or metabolic challenges during pregnancy, in which BBB disruption is also accompanied by changes in microglia number and activation^42,43^. These cells are crucial for vascular development, BBB function and integrity, and have been shown to be affected by altered microbial signaling^37,38,44^. However, in a model of CS, microglia numbers in the PVN have been shown to be decreased and in CA1 to be increased from birth to P3, indicating region- and time-dependent dynamic regulatory processes^45^. In parallel, studies in MIA models report transient peaks in microglial phagocytic activity in cortical regions such as the anterior cingulate cortex at P8–P10^17^, further highlighting the dynamic and context-specific nature of microglial phenotypes during early postnatal development. Together, these observations suggest that the increased brain tracer permeability observed here is unlikely to reflect a primary, microglia-driven BBB disruption.

Similarly to microglia, during embryonic development and early postnatal life, blood vessels undergo periods of intense plasticity characterized by changes in vessel permeability as well as network remodeling. This phenomenon is particularly heightened after the first postnatal week (P8 to P12), during which cortical blood flow increases as a consequence of an angiogenic burst^46^. In our study, CS did not have a strong effect on vessel network organization, but it did affect endothelial properties (increased PDX fluorescence in the 3V ChP) and permeability (increased PV1 fluorescence in the 3V ChP). PDX is a significant component of the endothelial glycocalyx and an essential component of barrier function^47,48^. In models of vascular remodeling and tumor-induced angiogenesis, degradation of the glycocalyx, alongside the tight junction network, strongly influenced microvascular permeability^49^. Thus, high PDX intensity observed in ChP vessels was consistent with altered glycocalyx organization, which could underlie increased vessel permeability. Additionally, PV1 is a protein associated with vasculature integrity and located in fenestrae, caveolae, and transendothelial channels of blood vessels. In physiological conditions, PV1 is only expressed during embryogenesis in the continuous barrier endothelium of the BBB^50^. The lack of fenestrae in the postnatal BBB allows for the tight restriction of exchanges from the blood to the brain parenchyma. As expected, our results highlighted that PV1 expression in the mPFC was overall very low and did not differ between groups. In contrast, vessels supplying the ChP are heavily fenestrated and classified as non-barrier endothelium. Studies investigating continuous fenestrated endothelia have shown that PV1 is necessary for vessel formation and that its absence leads to increased permeability of fenestrated vessels^51,52^. However, a recent study showed that gut inflammation (induced by dextran sodium sulfate-induced) led to the closure of the ChP endothelium^28^. This contradicts the classical idea that ChP fenestrated vessels are constantly permissive and part of the non-barrier endothelia. Here, we showed that CS led to increased PV1 intensity in the 3V ChP and concurrently to a widespread increase in tracer signal. Therefore, this central uptake of tracer could potentially reflect altered BBB function and/or tracer entry via periventricular interfaces, including the ChP endothelium. Notably, the most pronounced structural and immune alterations were observed at the BCSFB, supporting it as a key site of vulnerability in this model, particularly in the 3V ChP. Owing to their unique organization, localization and function, ChP vessels may represent an intermediary between continuous barrier and fenestrated non-barrier endothelia. Further studies should aim at characterizing ChP vessel dynamics upon modulation by extrinsic factors, specifically during the neonatal period.

Tight junctions represent determinant structural components of both brain barriers, as their network alteration is typically associated with impaired barrier function and neuroinflammation^30,39,40,53,54^. Although in our hands CS led to tracer leakage in the brain parenchyma, mRNA analysis showed only a trend towards a reduction of ZO-1 expression in the CS context and no difference in Ocln and Cldn5 expression between groups. Taken together, these results suggest that the tight junction network is preserved following CS and is unlikely to account for the increased barrier permeability observed in CS offspring. In the ChP, CS was not associated with overt disruption of the epithelial tight junction network but was accompanied by region-specific remodeling of this network, including increased maximum branch length and trends towards altered network complexity (number of endpoints) in the LV ChP. This suggests that the BCSFB epithelium may be dynamically regulated in response to CS-associated signals.

Furthermore, ChPs are also considered an immune barrier, protecting the brain from infections. In our CS model, we noted region-dependent changes in macrophage populations. In the LV ChP, an increase in the number of activated BAMs (Iba1^+^) and antigen-presenting cells was associated with a shift between BAM populations, indicating a reorganization of immune cells in response to CS. Such changes have previously been observed in the LV ChP in response to different inflammatory contexts^30,40^. In the 3V ChP, a decrease in the number of Iba1^+^ positive cells with phagocytic activity (Iba1^+^ CD68^+^) was observed. Recent work showed that macrophages in the LV ChP promoted barrier repair in inflammatory conditions^40^. Taken together, the increased maximum branch length of the tight junction network as well as the immune status of the ChPs after CS, suggest that similar barrier repair or protection mechanisms could be happening through immune and epithelial cells communication.

All in all, the tracer data indicates altered brain and 3V ChP tracer distribution but does not permit definitive localization of the site or route of barrier crossing (e.g., endothelial vs epithelial), as low-molecular weight tracers may rapidly diffuse or equilibrate within fenestrated vascular beds. Moreover, increased cortical parenchymal signals could arise from altered BBB function and/or from CSF-mediated routes following entry at periventricular interfaces.

In parallel with the observed alterations in brain barrier function, CS was associated with early changes in brain metabolic profiles. Given that delivery mode alters both gut permeability^19^ and microbial colonization during the first postnatal week^7^, and that barrier systems regulate the exchange of circulating metabolites with the brain^20^, we examined whether CS was accompanied by metabolic shifts in plasma and the developing cortex at P9. Lower NAA in the cortex of CS pups is noteworthy, as NAA is widely considered a marker of neuronal integrity, mitochondrial function, and neuron-glia metabolic coupling^55^. Previous work in neonatal piglets showed that brain NAA abundance was sensitive to systemic mediators such as cortisol^56^, raising the possibility that perinatal endocrine signals such as CS contribute to metabolic alterations. Together, these findings suggest that CS may also influence early neuronal metabolic capacity or delay aspects of neuron-glia interactions. Additionally, the observed decrease in malate, a key intermediate of the tricarboxylic acid (TCA) cycle and mitochondrial redox balance, may indicate a disruption of central energy metabolism^57^.

Finally, the increase in phosphocholine may point towards altered membrane phospholipid turnover or delayed membrane maturation. Phosphocholine is a precursor for phosphatidylcholine synthesis and is commonly associated with membrane remodeling during periods of rapid brain growth^58^. An imbalance between membrane synthesis and neuronal metabolic maturation could reflect asynchronous development of structural and metabolic components in the CS brain, potentially rendering specific regions such as the prefrontal cortex more vulnerable during early postnatal life.

Altogether, these metabolomic alterations indicate that CS delivery is associated with early changes in neuronal energy metabolism, amino acid utilization, and membrane dynamics in the cortex, paralleling the barrier vulnerability observed at this stage and supporting the notion that delivery mode shapes multiple interrelated aspects of early brain physiology.

Our data highlighted region-dependent differences induced by CS. Studies performed at later developmental timepoints have shown that CS affects inflammatory status around weaning and adolescence^59^ (Ratsika et al., unpublished observations). Here, we provide evidence that CS impacts the immune system as early as the end of the first postnatal week and that the effects of CS are complex and region dependent. CS differentially modulated the LV and 3V ChPs with vessels more affected by the delivery mode in the 3V ChP compared to the LV ChP. While the LV ChPs are closer to the cerebral cortex, the 3V ChP is situated near major neuroendocrine structures. This is associated with differences in structure and morphology, developmental patterns, functions, and sensitivity to modulations. Interestingly, the Hypothalamic-Pituitary-Adrenal (HPA) axis has been found to be dysregulated in infants born by CS^60^. In line with this, several preclinical CS studies have reported long-lasting alterations in hypothalamic neuroendocrine systems, including vasopressin- and oxytocin-related pathways, as well as early-life physiological stress responses and social behavior, highlighting the particular vulnerability of hypothalamic and periventricular regions to CS^6,61,62^. Moreover, we have also recently shown that CS delivery shifted the phase of microbial and endocrine rhythms at P9 (Gavioli et al., unpublished observations). In addition, preclinical studies have shown that early-life oxytocin administration attenuates CS-associated behavioral alterations and promotes sealing of both the BBB and BCSFB in mice with genetic vulnerability to neurodevelopmental disorders^63,64^. Thus, CS may affect the 3V ChP permeability through hormonal dysregulation. Interestingly, circumventricular organs are known to display dynamic fenestrated vessels reminiscent of ChP vessels, which are characterized by high permeability in early life. It is possible to infer that these vessels may also be affected by the delivery mode. This highlights the need to study LV, 3V, and 4V ChPs together and to further characterize the effects of CS in the hypothalamic area.

Our study bears some limitations. While we studied extensively qualitatively and quantitatively several proteins involved in barrier function and permeability, we could have benefited from transmission electron microscopy imaging to study the NVU integrity and cytoarchitecture to complement our results. In addition, CS pups required immediate placement with a foster dam at birth, whereas vaginally born controls remained with their biological dam. This paradigm differs from classical cross-fostering, which involves maternal reassignment after the establishment of dam-pup interaction. Although previous work using similar CS paradigms indicates that maternal reassignment alone does not reproduce CS-associated phenotypes^7^, we cannot entirely exclude subtle contributions from early postnatal environment or perinatal handling. Additionally, we have focused our investigation on a specific timepoint and brain regions, based on evidence of predisposition in the context of CS. Future studies should focus on the hypothalamus, fourth ventricle choroid plexus, and cerebellum and extend this investigation to later time points^59,65,66^.

Overall, our results point towards brain barrier impairment at the end of the first postnatal week, highlighting for the first time the determinant role of the BCSFB in the context of delivery mode and perinatal insults. While we did not directly investigate implications from the gut microbiota in this study, others have recently shown that specific bacterial taxa are associated with CS behavioral and inflammation changes in humans and mice^15,67–69^. Additionally, probiotics and prebiotics at P7 were shown to rescue selective behavioral alterations and gut barrier function in mice, strengthening the need to extend those gut microbiome interventions in the context of perinatal insults and brain barriers^7,19^. Similarly, we have shown that microbial-derived metabolites such as butyrate and propionate induced actin cytoskeletal rearrangement and protected BBB function^70^, and more importantly highlighted BCSFB alterations in germ free mice, i.e. mice completely depleted from a microbiome^71^. Taken together, these findings underscore the need to consider the BCSFB as a central and vulnerable interface during early-life adversity, and open new avenues for mechanistic exploration and therapeutic strategies aimed at safeguarding neurodevelopment.

## Supporting information

Turpin, Morael, Cergneux et al. 2026 SI

## RESOURCE AVAILABILITY

Requests for further information and resources should be directed to and will be fulfilled by the lead contact, María Rodriguez Aburto.

### Materials availability

- This study did not generate new unique reagents.

### Data and code availability

- Raw data reported in this paper will be shared by the lead contact upon request.
- Any additional information required to reanalyze the data reported in this paper is available from the lead contact upon request.
- This paper does not report original code.

## ACKNOWLEDGMENTS

M.R.A. was initially supported by a Marie Skłodowska-Curie Actions (MSCA) Individual Fellowship (NeuroVascOme, Grant Agreement 895999) and by the Marie Skłodowska-Curie Actions COFUND APC Postdoctoral Excellence (APEX; Grant Agreement 754535) Marie Skłodowska-Curie COFUND Fellowship, hosted by J.F.C. at University College Cork. This work was subsequently supported by a European Research Council (ERC) Starting Grant (RADIOGUT; Grant Agreement No. 101040951) and by a Science Foundation Ireland/Irish Research Council (SFI/IRC) Pathway Award (21/PATH-S/9424). V.T. and L.M. benefited from funding from the European Union’s Horizon 2020 research and innovation programme under the Marie Skłodowska-Curie grant agreement No 101034270. J.F.C. & G.C. are PIs in APC Microbiome Ireland, which was funded by Science Foundation Ireland (SFI/12/RC/2273_P2).

We thank Colette Manley, Patrick Fitzgerald, Gerard M. Moloney, Suzanne Crotty, and Tara Foley for excellent technical assistance. We thank Jatin Nagpal and Jerry Reen for providing access to and support with whole-brain imaging for tracer experiments. We thank Lily Keane, Austeja Baleviciute, and Tadhg Crowley for expert support with flow cytometry.

## AUTHOR CONTRIBUTIONS

M.R.A. and J.F.C conceived and designed the study, supervised the research, and secured funding. A.J.C.C., J.M., V.T., and M.R.A. planned and designed mouse experiments. A.J.C.C., J.M., V.T., H.B., L.M., E.G.K., A.R., K.G., P.V.S., J.S.H., performed mouse experiments. J.S., E.A., A.B. and N.J.L. performed metabolomics and analyzed data. A.J.C.C., J.M., V.T., H.B., E.C., C.T., A.M., analyzed data. A.J.C.C., J.M., V.T., H.B., J.F.C. and M.R.A. wrote the manuscript. All authors discussed the results and commented on the manuscript.

## FUNDING SOURCES

European Research Council, https://ror.org/0472cxd90, 101040951

European Union, 895999, 754535, 101034270

Science Foundation Ireland, 21/PATH-S/9424

## DECLARATION OF GENERATIVE AI AND AI-ASSISTED TECHNOLOGIES IN THE WRITING PROCESS

During the preparation of this work, the author(s) used Microsoft Copilot to refine manuscript wording. After using this tool or service, the author(s) reviewed and edited the content as needed and take full responsibility for the content of the publication.

## SUPPLEMENTAL INFORMATION

Document S1. Figures S1–S5, Table S1, and supplemental references.

## REFERENCES

1. Angolile, C.M., Max, B.L., Mushemba, J., and Mashauri, H.L. (2023). Global increased cesarean section rates and public health implications: A call to action. Health Sci. Rep. 6, e1274. 10.1002/hsr2.1274.

2. Boerma, T., Ronsmans, C., Melesse, D.Y., Barros, A.J.D., Barros, F.C., Juan, L., Moller, A.-B., Say, L., Hosseinpoor, A.R., Yi, M., et al. (2018). Global epidemiology of use of and disparities in caesarean sections. Lancet 392, 1341–1348. 10.1016/S0140-6736(18)31928-7.

3. Kristensen, K., and Henriksen, L. (2016). Cesarean section and disease associated with immune function. J. Allergy Clin. Immunol. 137, 587–590. 10.1016/j.jaci.2015.07.040.

4. Liu, X., Zhou, J., Chen, J., Li, L., Yuan, L., Li, S., Sun, X., and Zhou, X. (2024). Risk of asthma and allergies in children delivered by cesarean section: A comprehensive systematic review. J. Allergy Clin. Immunol. Pract. 12, 2764–2773. 10.1016/j.jaip.2024.06.022.

5. Sandall, J., Tribe, R.M., Avery, L., Mola, G., Visser, G.H., Homer, C.S., Gibbons, D., Kelly, N.M., Kennedy, H.P., Kidanto, H., et al. (2018). Short-term and long-term effects of caesarean section on the health of women and children. Lancet 392, 1349–1357. 10.1016/S0140-6736(18)31930-5.

6. Castillo-Ruiz, A., Mosley, M., Jacobs, A.J., Hoffiz, Y.C., and Forger, N.G. (2018). Birth delivery mode alters perinatal cell death in the mouse brain. Proc. Natl. Acad. Sci. U. S. A. 115, 11826–11831. 10.1073/pnas.1811962115.

7. Morais, L.H., Golubeva, A.V., Moloney, G.M., Moya-Pérez, A., Ventura-Silva, A.P., Arboleya, S., Bastiaanssen, T.F.S., O’Sullivan, O., Rea, K., Borre, Y., et al. (2020). Enduring Behavioral Effects Induced by Birth by Caesarean Section in the Mouse. Curr. Biol. 30, 3761-3774.e6. 10.1016/j.cub.2020.07.044.

8. Cabré, S., Ratsika, A., Rea, K., Stanton, C., and Cryan, J.F. (2022). Animal models for assessing impact of C-section delivery on biological systems. Neurosci. Biobehav. Rev. 135, 104555. 10.1016/j.neubiorev.2022.104555.

9. Zhang, T., Sidorchuk, A., Sevilla-Cermeño, L., Vilaplana-Pérez, A., Chang, Z., Larsson, H., Mataix-Cols, D., and Fernández de la Cruz, L. (2019). Association of cesarean delivery with risk of neurodevelopmental and psychiatric disorders in the offspring: A systematic review and meta-analysis. JAMA Netw. Open 2, e1910236. 10.1001/jamanetworkopen.2019.10236.

10. Lagercrantz, H., and Slotkin, T.A. (1986). The “stress” of being born. Sci. Am. 254, 100–107. 10.1038/scientificamerican0486-100.

11. Hillman, N.H., Kallapur, S.G., and Jobe, A.H. (2012). Physiology of transition from intrauterine to extrauterine life. Clin. Perinatol. 39, 769–783. 10.1016/j.clp.2012.09.009.

12. Kenkel, W. (2021). Birth signalling hormones and the developmental consequences of caesarean delivery. J. Neuroendocrinol. 33, e12912. 10.1111/jne.12912.

13. Dominguez-Bello, M.G., Costello, E.K., Contreras, M., Magris, M., Hidalgo, G., Fierer, N., and Knight, R. (2010). Delivery mode shapes the acquisition and structure of the initial microbiota across multiple body habitats in newborns. Proc. Natl. Acad. Sci. U. S. A. 107, 11971–11975. 10.1073/pnas.1002601107.

14. Mueller, N.T., Whyatt, R., Hoepner, L., Oberfield, S., Dominguez-Bello, M.G., Widen, E.M., Hassoun, A., Perera, F., and Rundle, A. (2015). Prenatal exposure to antibiotics, cesarean section and risk of childhood obesity. Int. J. Obes. (Lond) 39, 665–670. 10.1038/ijo.2014.180.

15. Shao, Y., Forster, S.C., Tsaliki, E., Vervier, K., Strang, A., Simpson, N., Kumar, N., Stares, M.D., Rodger, A., Brocklehurst, P., et al. (2019). Stunted microbiota and opportunistic pathogen colonization in caesarean-section birth. Nature 574, 117–121. 10.1038/s41586-019-1560-1.

16. Gour, A., Boergens, K.M., Heike, N., Hua, Y., Laserstein, P., Song, K., and Helmstaedter, M. (2021). Postnatal connectomic development of inhibition in mouse barrel cortex. Science 371, eabb4534. 10.1126/science.abb4534.

17. Block, C.L., Eroglu, O., Mague, S.D., Smith, C.J., Ceasrine, A.M., Sriworarat, C., Blount, C., Beben, K.A., Malacon, K.E., Ndubuizu, N., et al. (2022). Prenatal environmental stressors impair postnatal microglia function and adult behavior in males. Cell Rep. 40, 111161. 10.1016/j.celrep.2022.111161.

18. Ayata, P., Badimon, A., Strasburger, H.J., Duff, M.K., Montgomery, S.E., Loh, Y.-H.E., Ebert, A., Pimenova, A.A., Ramirez, B.R., Chan, A.T., et al. (2018). Epigenetic regulation of brain region-specific microglia clearance activity. Nat. Neurosci. 21, 1049–1060. 10.1038/s41593-018-0192-3.

19. Silva, A.P.V., Moloney, G.M., Sequeira, A.M., Liber, M., Bastiaanssen, T., Rea, K., Fitzgerald, P., Golubeva, A., Rodriguez-Aburto, M., Renes, I.B., et al. (2025). Targeting the Microbiota reverses C-section-induced effects on intestinal permeability, Microbiota composition, and amygdala gene expression in the mouse. Neurogastroenterol. Motil. 37, e70107. 10.1111/nmo.70107.

20. Aburto, M.R., and Cryan, J.F. (2024). Gastrointestinal and brain barriers: unlocking gates of communication across the microbiota-gut-brain axis. Nat. Rev. Gastroenterol. Hepatol. 21, 222–247. 10.1038/s41575-023-00890-0.

21. Dion-Albert, L., Dudek, K.A., Russo, S.J., Campbell, M., and Menard, C. (2023). Neurovascular adaptations modulating cognition, mood, and stress responses. Trends Neurosci. 10.1016/j.tins.2023.01.005.

22. Coelho-Santos, V., and Shih, A.Y. (2020). Postnatal development of cerebrovascular structure and the neurogliovascular unit. Wiley Interdiscip. Rev. Dev. Biol. 9, e363. 10.1002/wdev.363.

23. Paton, S.E.J., Solano, J.L., Coulombe-Rozon, F., Lebel, M., and Menard, C. (2023). Barrier-environment interactions along the gut-brain axis and their influence on cognition and behaviour throughout the lifespan. J. Psychiatry Neurosci. 48, E190–E208. 10.1503/jpn.220218.

24. Lehtinen, M.K., Zappaterra, M.W., Chen, X., Yang, Y.J., Hill, A.D., Lun, M., Maynard, T., Gonzalez, D., Kim, S., Ye, P., et al. (2011). The cerebrospinal fluid provides a proliferative niche for neural progenitor cells. Neuron 69, 893–905. 10.1016/j.neuron.2011.01.023.

25. Johansson, P.A. (2014). The choroid plexuses and their impact on developmental neurogenesis. Front. Neurosci. 8, 340. 10.3389/fnins.2014.00340.

26. Fame, R.M., and Lehtinen, M.K. (2020). Emergence and developmental roles of the cerebrospinal fluid system. Dev. Cell 52, 261–275. 10.1016/j.devcel.2020.01.027.

27. Bitanihirwe, B.K.Y., Lizano, P., and Woo, T.-U.W. (2022). Deconstructing the functional neuroanatomy of the choroid plexus: an ontogenetic perspective for studying neurodevelopmental and neuropsychiatric disorders. Mol. Psychiatry. 10.1038/s41380-022-01623-6.

28. Carloni, S., Bertocchi, A., Mancinelli, S., Bellini, M., Erreni, M., Borreca, A., Braga, D., Giugliano, S., Mozzarelli, A.M., Manganaro, D., et al. (2021). Identification of a choroid plexus vascular barrier closing during intestinal inflammation. Science 374, 439–448. 10.1126/science.abc6108.

29. Saunders, N.R., Dziegielewska, K.M., Fame, R.M., Lehtinen, M.K., and Liddelow, S.A. (2023). The choroid plexus: a missing link in our understanding of brain development and function. Physiol. Rev. 103, 919–956. 10.1152/physrev.00060.2021.

30. Cui, J., Shipley, F.B., Shannon, M.L., Alturkistani, O., Dani, N., Webb, M.D., Sugden, A.U., Andermann, M.L., and Lehtinen, M.K. (2020). Inflammation of the Embryonic Choroid Plexus Barrier following Maternal Immune Activation. Dev. Cell 55, 617-628.e6. 10.1016/j.devcel.2020.09.020.

31. Cui, J., Xu, H., and Lehtinen, M.K. (2021). Macrophages on the margin: choroid plexus immune responses. Trends Neurosci. 44, 864–875. 10.1016/j.tins.2021.07.002.

32. Segarra, M., Aburto, M.R., and Acker-Palmer, A. (2021). Blood-brain barrier dynamics to maintain brain homeostasis. Trends Neurosci. 44, 393–405. 10.1016/j.tins.2020.12.002.

33. Sweeney, M.D., Zhao, Z., Montagne, A., Nelson, A.R., and Zlokovic, B.V. (2019). Blood-brain barrier: From physiology to disease and back. Physiol. Rev. 99, 21–78. 10.1152/physrev.00050.2017.

34. Ben-Zvi, A., Lacoste, B., Kur, E., Andreone, B.J., Mayshar, Y., Yan, H., and Gu, C. (2014). Mfsd2a is critical for the formation and function of the blood-brain barrier. Nature 509, 507–511. 10.1038/nature13324.

35. Liddelow, S.A., Temple, S., Møllgård, K., Gehwolf, R., Wagner, A., Bauer, H., Bauer, H.-C., Phoenix, T.N., Dziegielewska, K.M., and Saunders, N.R. (2012). Molecular characterisation of transport mechanisms at the developing mouse blood-CSF interface: a transcriptome approach. PLoS One 7, e33554. 10.1371/journal.pone.0033554.

36. Llaó-Cid, C., Peguera, B., Kobialka, P., Decker, L., Vogenstahl, J., Alivodej, N., Srivastava, S., Jin, J., Kirchmaier, B.C., Milla, C., et al. (2024). Vascular FLRT2 regulates venous-mediated angiogenic expansion and CNS barriergenesis. Nat. Commun. 15, 10372. 10.1038/s41467-024-54570-x.

37. Chen, S., Li, J., Meng, S., He, T., Shi, Z., Wang, C., Wang, Y., Cao, H., Huang, Y., Zhang, Y., et al. (2023). Microglia and macrophages in the neuro-glia-vascular unit: From identity to functions. Neurobiol. Dis. 179, 106066. 10.1016/j.nbd.2023.106066.

38. Bijnen, M., Sridhar, S., Keller, A., and Greter, M. (2025). Brain macrophages in vascular health and dysfunction. Trends Immunol. 46, 46–60. 10.1016/j.it.2024.11.012.

39. Xie, J., Bruggeman, A., De Nolf, C., Vandendriessche, C., Van Imschoot, G., Van Wonterghem, E., Vereecke, L., and Vandenbroucke, R.E. (2023). Gut microbiota regulates blood-cerebrospinal fluid barrier function and Aβ pathology. EMBO J. 42, e111515. 10.15252/embj.2022111515.

40. Xu, H., Lotfy, P., Gelb, S., Pragana, A., Hehnly, C., Byer, L.I.J., Shipley, F.B., Zawadzki, M.E., Cui, J., Deng, L., et al. (2024). The choroid plexus synergizes with immune cells during neuroinflammation. Cell 187, 4946-4963.e17. 10.1016/j.cell.2024.07.002.

41. Dominguez-Belloso, A., Schmutz, S., Novault, S., Travier, L., and Deczkowska, A. (2022). Isolation and characterization of the immune cells from micro-dissected mouse choroid plexuses. J. Vis. Exp. 10.3791/63487.

42. Zhao, Q., Dai, W., Chen, H.Y., Jacobs, R.E., Zlokovic, B.V., Lund, B.T., Montagne, A., and Bonnin, A. (2022). Prenatal disruption of blood-brain barrier formation via cyclooxygenase activation leads to lifelong brain inflammation. Proc. Natl. Acad. Sci. U. S. A. 119, e2113310119. 10.1073/pnas.2113310119.

43. Thion, M.S., Low, D., Silvin, A., Chen, J., Grisel, P., Schulte-Schrepping, J., Blecher, R., Ulas, T., Squarzoni, P., Hoeffel, G., et al. (2018). Microbiome influences prenatal and adult microglia in a sex-specific manner. Cell 172, 500-516.e16. 10.1016/j.cell.2017.11.042.

44. Blair, H.J., Morales, L., Cryan, J.F., and Aburto, M.R. (2025). Neuroglia and the microbiota-gut-brain axis. Handb. Clin. Neurol. 209, 171–196. 10.1016/B978-0-443-19104-6.00001-2.

45. Castillo-Ruiz, A., Cisternas, C.D., Sturgeon, H., and Forger, N.G. (2022). Birth triggers an inflammatory response in the neonatal periphery and brain. Brain Behav. Immun. 104, 122–136. 10.1016/j.bbi.2022.05.021.

46. Coelho-Santos, V., Berthiaume, A.-A., Ornelas, S., Stuhlmann, H., and Shih, A.Y. (2021). Imaging the construction of capillary networks in the neonatal mouse brain. Proc. Natl. Acad. Sci. U. S. A. 118, e2100866118. 10.1073/pnas.2100866118.

47. Reitsma, S., Slaaf, D.W., Vink, H., van Zandvoort, M.A.M.J., and oude Egbrink, M.G.A. (2007). The endothelial glycocalyx: composition, functions, and visualization. Pflugers Arch. 454, 345–359. 10.1007/s00424-007-0212-8.

48. Kutuzov, N., Flyvbjerg, H., and Lauritzen, M. (2018). Contributions of the glycocalyx, endothelium, and extravascular compartment to the blood-brain barrier. Proc. Natl. Acad. Sci. U. S. A. 115, E9429–E9438. 10.1073/pnas.1802155115.

49. Zeng, Y., and Fu, B.M. (2025). Angiogenesis and microvascular permeability. Cold Spring Harb. Perspect. Med. 15, a041163. 10.1101/cshperspect.a041163.

50. Hallmann, R., Mayer, D.N., Berg, E.L., Broermann, R., and Butcher, E.C. (1995). Novel mouse endothelial cell surface marker is suppressed during differentiation of the blood brain barrier. Dev. Dyn. 202, 325–332. 10.1002/aja.1002020402.

51. Herrnberger, L., Seitz, R., Kuespert, S., Bösl, M.R., Fuchshofer, R., and Tamm, E.R. (2012). Lack of endothelial diaphragms in fenestrae and caveolae of mutant Plvap-deficient mice. Histochem. Cell Biol. 138, 709–724. 10.1007/s00418-012-0987-3.

52. Stan, R.V., Tse, D., Deharvengt, S.J., Smits, N.C., Xu, Y., Luciano, M.R., McGarry, C.L., Buitendijk, M., Nemani, K.V., Elgueta, R., et al. (2012). The diaphragms of fenestrated endothelia: gatekeepers of vascular permeability and blood composition. Dev. Cell 23, 1203–1218. 10.1016/j.devcel.2012.11.003.

53. Luissint, A.-C., Artus, C., Glacial, F., Ganeshamoorthy, K., and Couraud, P.-O. (2012). Tight junctions at the blood brain barrier: physiological architecture and disease-associated dysregulation. Fluids Barriers CNS 9, 23. 10.1186/2045-8118-9-23.

54. Greene, C., Hanley, N., and Campbell, M. (2020). Blood-brain barrier associated tight junction disruption is a hallmark feature of major psychiatric disorders. Transl. Psychiatry 10, 373. 10.1038/s41398-020-01054-3.

55. Moffett, J.R., Ross, B., Arun, P., Madhavarao, C.N., and Namboodiri, A.M.A. (2007). N-Acetylaspartate in the CNS: from neurodiagnostics to neurobiology. Prog. Neurobiol. 81, 89–131. 10.1016/j.pneurobio.2006.12.003.

56. Mudd, A.T., Berding, K., Wang, M., Donovan, S.M., and Dilger, R.N. (2017). Serum cortisol mediates the relationship between fecal Ruminococcus and brain N-acetylaspartate in the young pig. Gut Microbes 8, 589–600. 10.1080/19490976.2017.1353849.

57. Martínez-Reyes, I., and Chandel, N.S. (2020). Mitochondrial TCA cycle metabolites control physiology and disease. Nat. Commun. 11, 102. 10.1038/s41467-019-13668-3.

58. Marcucci, H., Paoletti, L., Jackowski, S., and Banchio, C. (2010). Phosphatidylcholine biosynthesis during neuronal differentiation and its role in cell fate determination. J. Biol. Chem. 285, 25382–25393. 10.1074/jbc.M110.139477.

59. Di Miceli, M., Rossitto, M., Martinat, M., Marchaland, F., Kharbouche, S., Graland, M., Younes, F., Séré, A., Aubert, A., Khabbaz, L.R., et al. (2024). Modified neuroimmune processes and emotional behaviour in weaned and late adolescent male and female mice born via caesarean section. Sci. Rep. 14, 29807. 10.1038/s41598-024-80770-y.

60. Martinez, L.D., Glynn, L.M., Sandman, C.A., Wing, D.A., and Davis, E.P. (2020). Cesarean delivery and infant cortisol regulation. Psychoneuroendocrinology 122, 104862. 10.1016/j.psyneuen.2020.104862.

61. Ramlall, E.K., Hall, M.A.L., Forger, N.G., and Castillo-Ruiz, A. (2021). Cesarean birth elicits long-term effects on vasopressin and oxytocin neurons in the hypothalamic paraventricular nucleus of mice. Horm. Behav. 136, 105080. 10.1016/j.yhbeh.2021.105080.

62. Hoffiz, Y.C., Castillo-Ruiz, A., Hall, M.A.L., Hite, T.A., Gray, J.M., Cisternas, C.D., Cortes, L.R., Jacobs, A.J., and Forger, N.G. (2021). Birth elicits a conserved neuroendocrine response with implications for perinatal osmoregulation and neuronal cell death. Sci. Rep. 11, 2335. 10.1038/s41598-021-81511-1.

63. Morais, L.H., Golubeva, A.V., Casey, S., Scott, K.A., Ramos Costa, A.P., Moloney, G.M., Dinan, T.G., and Cryan, J.F. (2021). Early-life oxytocin attenuates the social deficits induced by caesarean-section delivery in the mouse. Neuropsychopharmacology 46, 1958–1968. 10.1038/s41386-021-01040-3.

64. Castellani, G., Ciampoli, M., Benedetti, A., Ferretti, V., Trigilio, G., Barcik, W., Busnelli, M., Contarini, G., Paolini, C., Devroye, C., et al. (2025). Oxytocin seals the blood-brain barrier, improving 22q11.2 deletion syndrome trajectories. Brain, awaf112. 10.1093/brain/awaf112.

65. Chiesa, M., Guimond, D., Tyzio, R., Pons-Bennaceur, A., Lozovaya, N., Burnashev, N., Ferrari, D.C., and Ben-Ari, Y. (2019). Term or preterm cesarean section delivery does not lead to long-term detrimental consequences in mice. Cereb. Cortex 29, 2424–2436. 10.1093/cercor/bhy112.

66. Chiesa, M., Ferrari, D.C., and Ben-Ari, Y. (2020). Alteration in the time and/or mode of delivery differentially modulates early development in mice. Mol. Brain 13, 34. 10.1186/s13041-020-00578-5.

67. Stewart, C.J., Ajami, N.J., O’Brien, J.L., Hutchinson, D.S., Smith, D.P., Wong, M.C., Ross, M.C., Lloyd, R.E., Doddapaneni, H., Metcalf, G.A., et al. (2018). Temporal development of the gut microbiome in early childhood from the TEDDY study. Nature 562, 583–588. 10.1038/s41586-018-0617-x.

68. Barone, M., Ramayo-Caldas, Y., Estellé, J., Tambosco, K., Chadi, S., Maillard, F., Gallopin, M., Planchais, J., Chain, F., Kropp, C., et al. (2023). Gut barrier-microbiota imbalances in early life lead to higher sensitivity to inflammation in a murine model of C-section delivery. Microbiome 11, 140. 10.1186/s40168-023-01584-0.

69. Morin, C., Faure, F., Mollet, J., Guenoun, D., Heydari-Olya, A., Sautet, I., Diao, S., Faivre, V., Pansiot, J., Tabet, L., et al. (2025). C-section and systemic inflammation synergize to disrupt the neonatal gut microbiota and brain development in a model of prematurity. Brain Behav. Immun. 123, 824–837. 10.1016/j.bbi.2024.10.023.

70. Knox, E.G., Aburto, M.R., Tessier, C., Nagpal, J., Clarke, G., O’Driscoll, C.M., and Cryan, J.F. (2022). Microbial-derived metabolites induce actin cytoskeletal rearrangement and protect blood-brain barrier function. iScience 25, 105648. 10.1016/j.isci.2022.105648.

71. Knox, E.G., Lynch, C.M.K., Lee, Y.S., O’Driscoll, C.M., Clarke, G., Cryan, J.F., and Aburto, M.R. (2023). The gut microbiota is important for the maintenance of blood-cerebrospinal fluid barrier integrity. Eur. J. Neurosci. 57, 233–241. 10.1111/ejn.15878.

