## Supplementary material for "Birth by Cesarean Section Remodels Neonatal Brain Barriers and Associated Immune and Metabolic Pathways in Mice": Turpin, Morael, Cergneux et al. 2026 SI

### 1. MATERIALS AND METHODS

#### 1.1. *Animal Husbandry*

NIH Swiss (HSD:NIHS) female and male breeders were purchased from Envigo, UK, and let to acclimatize for up to two weeks. For this study, outbred mice were preferred because of their highly reliable maternal behavior[1]. Mice were fed *ad libitum* with a conventional diet (2018S, Harlan Laboratories) and housed individually in a 12 hour day/night light cycle (lights on from 7:30 am to 7:30 pm) in 33x15x13cm white cages with environmental enrichment (bedding, nesting material, cylindrical tube).

#### 1.2. *Cesarean section*

At day 19.5 (20 days ±1 day), pregnant females were euthanized by cervical dislocation, and after disinfection with 70% ethanol, an incision was made in the abdominal cavity to expose the uterus. Pups were extracted one by one and placed on a sterile gauze on top of a heating pad. Sterile cotton buds were then used to massage their abdomen until spontaneous breathing was observed. Pups were then given to a foster dam who had vaginally delivered within the last 24 hours, and pups were rolled on bedding material from foster cages to encourage acceptance by the foster dam.

#### 1.3. *In vivo procedures, sample collection and tissue processing*

*Flow cytometry of the choroid plexus.* To lyse the ChP tissue, enzymatic digestion was first performed using collagenase type IV (17104019, Thermofisher) (400U per mL of PBS) for 1h at 37 °C with mild agitation. After one hour, flow staining (FACS) buffer (00-4222-26, eBioscience) was added to stop enzymatic activity. The cells were then washed twice with FACS buffer and stained with Zombie™ NIR 1:1000 (in PBS, 15min in the dark, 423105, Biolegends) for cell viability assessment. After washing, FcR blocking reagent (130-092-575, Miltenyi) and brilliant violet staining buffer (563794, BD Biosciences) were added for 10 min to increase the specificity of the immunofluorescent staining. The samples were then stained for 30 minutes in the dark with immune-specific conjugated monoclonal antibodies (SI Appendix Table S1).

Cells were washed and fixed with 4% ice-cold PFA (for 45min in the dark) and kept at 4 °C until further processing. Cells were analyzed on a BD FACSCelesta™ Cell Analyzer (BD Biosciences). Unstained and single-stained samples from spleen tissues were used to calculate the compensation matrix and accurately identify the populations of interest.

The gating strategies reported in the figures were generated by FlowJo™11 software (BD Biosciences) and adapted from Dominguez-Belloso et al., 2022[2]. Cells were gated on forward and side scatter properties followed by gating for live, single cells and immune cells (CD45$^+$). Border-associated macrophages (BAM) were divided into two different populations according to their expression of MHC-II: IA-IE$^+$ BAM (CD45$^+$ CD11b$^+$ F4/80$^+$ CX3CR1$^+$ IA-IE$^+$) and IA-IE- BAM (CD45$^+$ CD11b$^+$ F4/80$^+$ CX3CR1$^+$ IA-IE$^-$). The latter group includes Kolmer's epiplexus macrophages (CD11bhigh F4/80intermediate). Flow cytometry outputs included percentage of parent populations, and median fluorescence intensity (MFI) for activation of cell surface markers.

*Cryostat sectioning and immunostaining.* Brains were acclimatized to 4 °C for 24 hours, placed in a 30% sucrose-PBS solution (overnight, 4 °C), then in a 30% sucrose-PBS:OCT (1:1) solution (4583 Tissue-Tek, Sakura) (overnight, 4 °C), and finally in OCT (1h, 4 °C). Brains were then frozen in OCT at ultra-low temperature using dry-ice-chilled isopentane and kept at -80 °C for long-term storage. For immunostaining, slides were thawed at room temperature (RT) and washed with PBS.

Choroid plexus sections stained for Zonula Occludens 1 (ZO-1) and Plasmalemma Vesicle-Associated Protein 1 (PV1) were subjected to enzymatic antigen retrieval. Slides were incubated with Proteinase XIV (0.2 mg/mL in 0.05 mM Tris-buffered saline (TBS), pH7.4) for 45 s at 37°C (P5147, Sigma Aldrich). Slides were washed once with Tris 0.01 M and twice with PBS prior to permeabilization (PBS 0.3% Triton™ X-100, 10% NDS, 2% bovine serum albumin (BSA)) for 120 minutes at room temperature.

Slides with choroid plexus and prefrontal cortex regions were selected and incubated with primary antibodies (diluted in 0.1% PBS Tween®20 and 5% serum) overnight at 4°C. The following antibodies were used: rat anti-PV1 1:50 (550563, BD Biosciences), goat anti-podocalyxin 1:150 (AF1556, R&D), rabbit anti-CD68 1:150 (PA5-78996, Invitrogen), rabbit anti-ZO-1 1:150 (61-7300, Invitrogen), rabbit anti-Aquaporin 4 1:200 (AB2218, Millipore), rabbit anti-IBA1 1:500 (019-19741, Fujifilm), guinea pig anti-IBA1 1:500 (234-308, Synaptic Systems). Slides were then washed 3 times with PBS 0.1% Tween®20 for 10 minutes and incubated with secondary antibodies in PBS 0.1% Tween®20 and 2% NDS for 120 minutes. The following secondary antibodies conjugated to Alexa Fluor™ Plus −488, −555, −568, −594, and −647 were used: donkey anti-mouse/rabbit/goat/rat (Invitrogen) and guinea pig (706-605-148, Jackson Immuno Research) 1:200. Finally, slides were washed 3 times with PBS 0.1% Tween®20 for 10 minutes. Slides were mounted with coverslips (631-0137, VWR) using ProLong® Gold (P36930, Invitrogen) mounting medium and allowed to dry at room temperature.

##### *1.4. Image acquisition and figure layout*

Images were acquired using a laser scanning confocal microscope (FLUOVIEW FV3000, Olympus). Brightness and contrast of the figure images were adjusted using ImageJ/Fiji. Figures were prepared using Inkscape v1.4.2.

##### *1.5. Confocal image analyses*

###### *1.5.1. Vasculature analysis in the prefrontal cortex and choroid plexus.*

Prefrontal cortex and choroid plexus vessels labelled with IB4 and PDX respectively, were imaged at a magnification of 20X (5 stacks of 2µm and 16 stacks of 0.96µm respectively). Images were pre-processed using ImageJ to create a mask of the vasculature which was then analyzed with REAVER (MatLab, R2024a) automatically (0.621µm/px and 0.3107µm/px respectively) then manually curated. Each animal (n=1) represents an average of values obtained from 1-5 images. Vessel length, branchpoint counts, segment counts, and valency were normalized to individual choroid plexus areas.

###### *1.5.2. Tight junction analysis on the prefrontal cortex and choroid plexus.*

Images were taken at 20X magnification (z-stack=5, step size=2µm) and processed with ImageJ/Fiji. Automatic thresholding was applied on ZO-1, using the Otsu method. Vessel masks created as described previously were applied to the ZO-1 channel and the signal confined within the vascular ROI was measured. Quantitative parameters, including the area, perimeter, and area fraction, were compiled. The principal outcome measure used for statistical analysis was the area fraction representing tight junction coverage within the vascular compartment. Results from 4 to 6 images were combined to yield a single value per animal.
Additionally, analysis of tight junction ultrastructure was performed on images taken at 60X magnification (5 stacks, step size=2µm). Five tight junction (TJ) categories were determined by an experienced researcher: mess, tremble, twist, discontinuity, and amalgam. Results from 4 to 6 images were combined to yield a single value per animal.
Analysis of the choroid plexus epithelium included quantification of ZO-1 fluorescence at the cell-cell junction, and analysis of its network within the tissue. Each image was taken at 60X magnification and consisted of 3 stacks for the LV choroid plexuses and 5 stacks for the third ventricle (3V) choroid plexus (step size=0.96µm).

To quantify tight junction linearity, three cells per image were analyzed. For each tight junction branch, two measurements of length were reported:

- The length of a straight line going from one node of the branch to another
- The segmented length obtained by measuring the length of the tight junction branch considering twists and angles. This represents the true length of the tight junctions.

The linearity index is then obtained by dividing segmented length by straight line length. Tight junction branches with a linearity index close to 1 generally display less tight junction ruffles and are more linear.

To quantify ZO-1 fluorescence and analyze its network, a region of interest (ROI) was selected within the epithelium in each image and was then cropped. The background was then processed with Substract Background (Rolling Ball Radius=2 pixels), Remove Outliers (Radius=4 pixels, Threshold=1, which outliers=Bright), Median (radius=2 pixels), Enhance Local Contrast (CLAHE) (Blocksize=127, Histogram=256, Maximum Slope=3, Mask=none), and was binarized.

For the ZO-1 fluorescence quantification, the binarized image was then processed through the Analyze Particles function (Size=0.25-infinity, Circularity=0-1, Show=Masks). This created a mask of the tight junction network without background. The colors were then inverted, and a selection of the particles only (i.e. the tight junction network) was created and applied to the original picture of the ZO-1 fluorescence (raw ROI with an unprocessed background). The mean grey value was then reported for the selection. This output represented the intensity of the tight junction protein at the cell-cell junction.

To assess the complexity of the epithelial tight junction network, the binary images were skeletonized and the network analyzed using the neuroanatomy plugin. All skeleton outputs were normalized to the ROI area, excluding the mean branch length.

Finally, by ticking Summarize in the Analyze Particles window, a summary of the particle analysis was created and represented a further assessment of the ZO-1 network organization.

Each animal (n=1) represents an average of values obtained from 1-6 images with 1-2 ROI per image.

##### ***1.5.3. Macrophage analysis in the choroid plexus.***

Macrophages, labelled with Iba1 and CD68, were imaged at a magnification of 20X (z-stack=16, step size=1.5µm). Macrophage analysis was performed using ImageJ/Fiji. For each image, an outline of the ChP was traced, and the number of Iba1$^+$, CD68$^+$, and Iba1$^+$ CD68$^+$ cells were reported. Epiplexus macrophages were considered as macrophages on the margin of the ChP outline or just outside of the tissue. Each cell count was normalized to the respective ChP area, and the percentage of CD68+ cells within the Iba1$^+$ macrophage population was calculated from normalized values. Each animal (n=1) represents an average of values obtained from 1-4 images.

##### ***1.5.4. Neuro-glia-vascular components in prefrontal cortex and choroid plexus***

*(Cadaverine-555, Iba1, PDCX, PV1, AQP4 quantification).*

<u>*Tracer analysis.*</u> Cadaverine leakage was quantified by measuring mean fluorescence intensity with ImageJ/Fiji (NIH, Bethesda, MD, USA)[3]. For all regions (cortex, cerebellum, hypothalamus), one region of interest (ROI) was selected per hemisphere, and the mean grey values from both ROIs were averaged to yield a single representative value (n=1).

For high-resolution analyses, cadaverine fluorescence was further imaged at 20X magnification using z-stacks (choroid plexus: z-stack=16, step size=0.96µm; prefrontal cortex sections: 5 stacks, step size=2µm) and processed in ImageJ/Fiji. For ChP samples, ROIs were defined within the stroma or vascular structures using a PDX staining. For brain sections, ROIs were selected within the mPFC parenchyma. For each animal (n=1), fluorescence intensity corresponds to the mean grey value averaged from 1–6 images (2–3 ROIs per image) for the ChP and from 4–6 images for brain sections.

<u>*Microglia and vasculature analysis.*</u> Coronal brain sections containing the prefrontal cortex were used. Microglia and vessels, labelled with Iba1 and PDX respectively, were imaged at a magnification of 20X. For each animal, 3-8 images were acquired (z-stack=5, step size=2µm) maintaining constant acquisition settings (laser power, gain, and exposure time) across all samples to ensure comparability. These images were processed on ImageJ/Fiji. Iba1$^+$ and Iba1$^+$ PDX$^+$ were counted manually to identify microglia and vascular associated microglia (VAM). Iba1+ cell count was normalized to area, and VAM count was expressed as a percentage of all Iba1$^+$ cells (microglia). Iba1 fluorescence intensity was measured within microglia area occupancy.

Microglial morphology was analyzed using CellProfiler (version 4.2.8). Iba1$^+$ cells were segmented using 'IdentifyPrimaryObjects' module, skeletonized with 'MorphologicalSkeleton' module, and quantified using 'MeasureImageSkeleton' module. Branching density was calculated as the number of skeleton branches normalized to cell area. Cell compactness (perimeter²/area-based measure of shape irregularity) was obtained from 'MeasureObjectSizeShape'. Measurements were averaged per animal for downstream statistical analysis.

*Astrocytic endfeet and vasculature analysis.* Coronal brain sections containing the prefrontal cortex were used. Astrocytic end-feet and vessels, labelled with AQP4 and PDX respectively, were imaged at a magnification of 40X. For each animal, 6 images were acquired (z-stacks=6, step size=1 µm) maintaining constant acquisition settings (laser power, gain, and exposure time) across all samples to ensure comparability.

These images were processed and analyzed using a custom macro in ImageJ/Fiji. Briefly, a maximum intensity of Z-projection was generated for each channel. Blood vessels were automatically segmented from the PDX channel. The resulting vascular mask was refined (fill holes, dilate) to generate an expanded perivascular mask. This mask was applied to the APQ4 channel to isolate the perivascular AQP4 signal. The co-localization of AQP4 and PDX signals was determined using the Image Calculator process. For each individual vessel (10-18 per image), the following parameters were quantified: the area (in pixels) of PDX and APQ4, the mean intensity of AQP4 signal and the area of AQP4-PDX co-localization. From these measurements, the AQP4 coverage ratio (AQP4 area / PDX area) and the AQP4 thickness ratio [(AQP4 area – co-localization area) / PDX area] were calculated. Additionally, vessel morphology metrics, including area, perimeter and circularity were measured for each segmented vessel.

*Fenestrated vessels and vasculature analysis.* PV1 and PDX were imaged at a magnification of 60X and 20X respectively (z-stack=16 images, step size=0.96µm). ROIs were selected within the vessel network, and the mean grey value was reported for each selection. For PV1, choroid plexus vessels were identified using PDX (Alexa Fluor 405). Each animal (n=1) represents an average of values obtained from 1-6 images with 1-3 ROIs per image.

#### 2. FIGURES

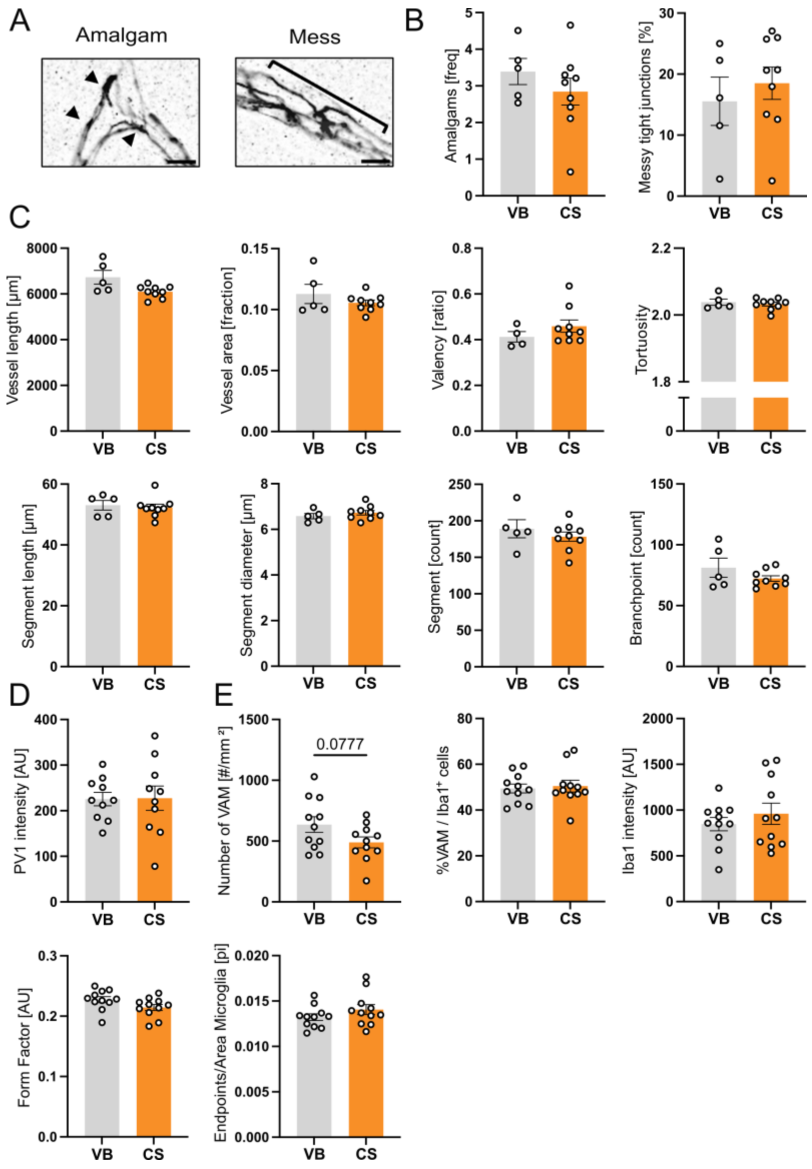


Fig. S1. Mode of delivery does not alter BBB components. (A) Representative confocal images of tight junction ultrastructure, visualized by ZO-1 immunostaining in cortical microvessels (Scale bar = 5 µm). (B) Analysis of tight junction ultrastructure characteristics: percentage of messy TJ in the vessels and frequency of amalgam in the TJ network. (mean ± SEM; t-test, n=5-9). (C) Quantitative analysis of the vascular network: vessel length, vessel area, valency, tortuosity, segment length, segment diameter, number of segments, number of branchpoint. (D) Fluorescence quantification of the vascular fenestration marker PV1 (mean ± SEM; Welch's t-test, n=10). (E) Quantitative and morphological analysis of microglial: vascular associated microglia (VAM) density, percentage of VAM among microglia population, Iba1 marker fluorescence intensity, form factor, branching density normalized to cell area (mean ± SEM; t-test, n=11).

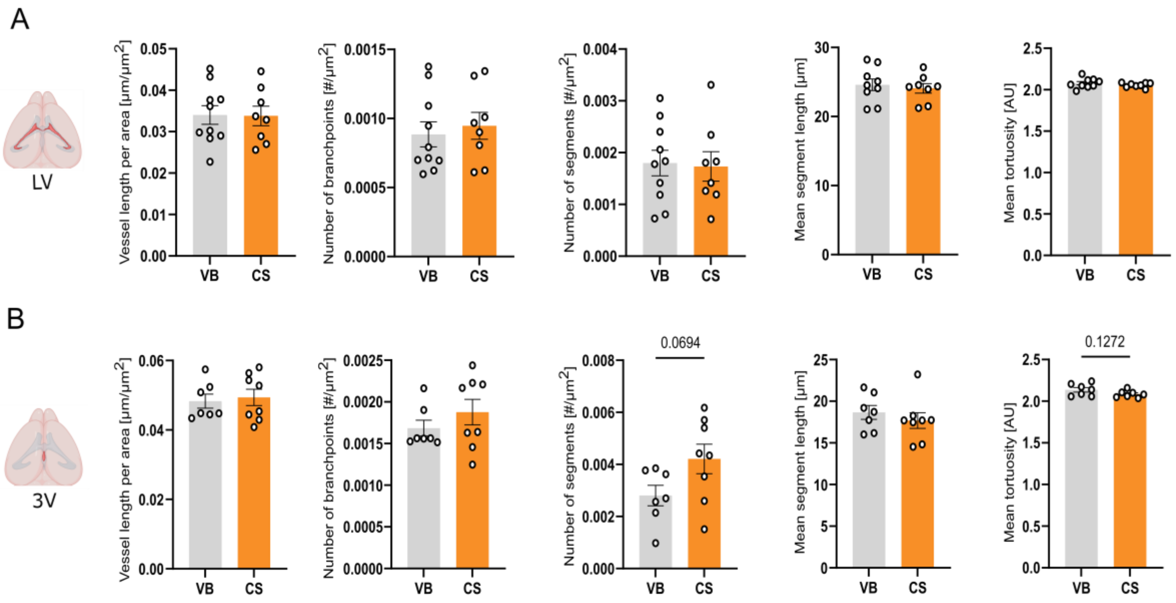


Fig. S2. Mode of delivery does not alter BCSFB vessel network. Leftmost panels: Vessel length normalized to ChP area. Left middle panels: Number of branchpoints normalized to ChP area. Middle panels: Number of segments normalized to ChP area. Right middle panels: Mean segment length. Rightmost panels: Mean vessel tortuosity. Upper panels (A): LV ChP (mean ± SEM; t-test, n=8-10); bottom panels (B): 3V ChP (mean ± SEM; t-test, n=7-8).

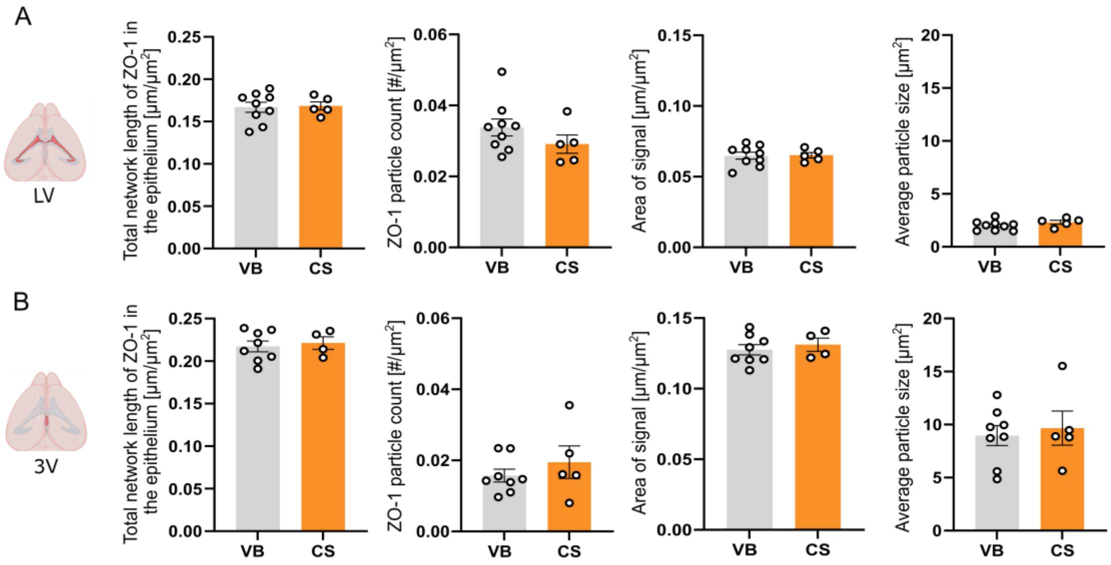


Fig. S3. Mode of delivery does not affect epithelial tight junction particles. Leftmost panels: Quantification of maximum branch length of the tight junction epithelial network. Left middle panels: Number of particles of ZO-1 fluorescence normalized to the overall area of the ChP epithelium. Right middle panels: Area of ZO-1 particles normalized to the overall area of the ChP epithelium. Rightmost panels: Average of the size of the ZO-1 particles. Upper panels (A): LV ChP (mean ± SEM; t-test, n=5-9); bottom panels (B): 3V ChP (mean ± SEM; t-test, n=4-8).

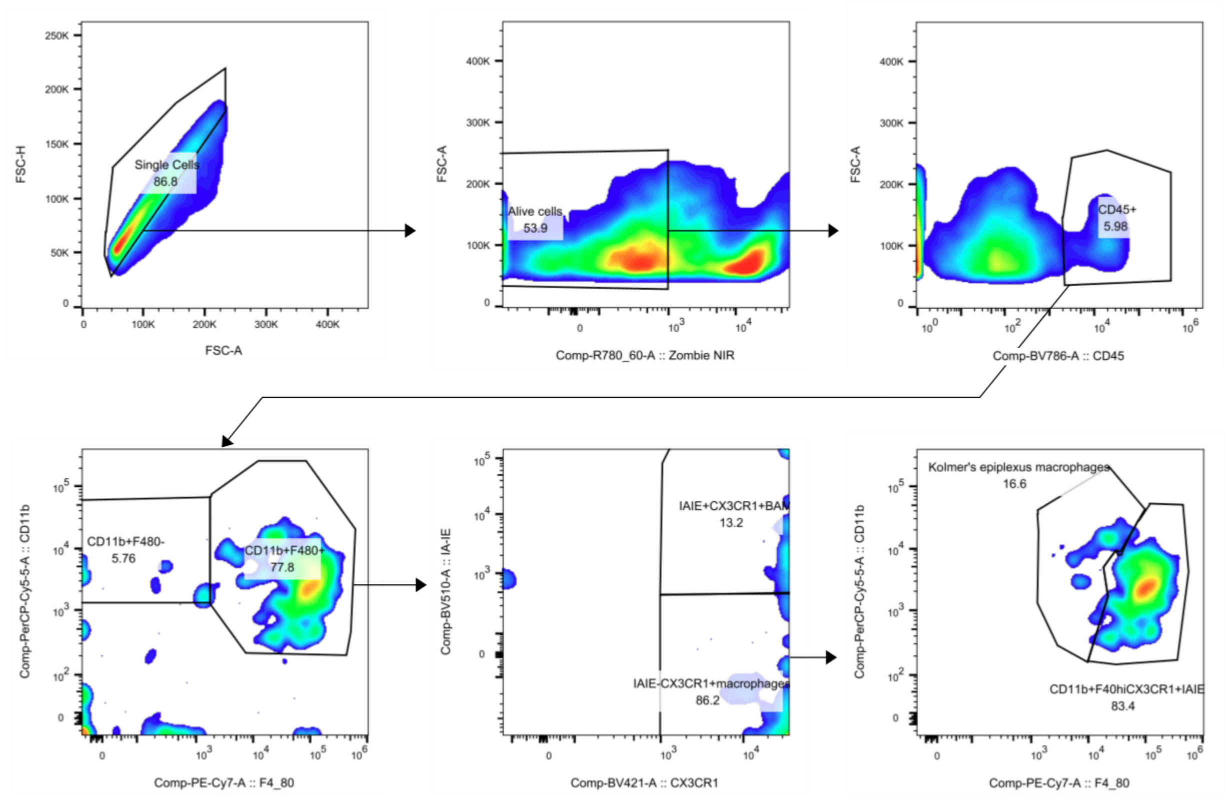


Fig. S4. Distribution of choroid plexus immune cells after flow cytometry. Upper panels: cells were gated on forward and side scatter properties followed by gating for live, single cells and immune cells ($CD45^{+}$). Border-associated macrophages (BAM) (Lower panel – left: $CD11b^{+}$ $F4/80^{+}$) were divided into two different populations according to their expression of MHC-II (Lower panel – middle): $IA\text{-}IE^{+}$ BAM ($CD45^{+}$ $CD11b^{+}$ $F4/80^{+}$ $CX3CR1^{+}$ $IA\text{-}IE^{+}$) and IA-IE- BAM ($CD45^{+}$ $CD11b^{+}$ $F4/80^{+}$ $CX3CR1^{+}$ $IA\text{-}IE^{-}$). The latter group includes Kolmer's epiplexus macrophages (Lower panel – right): CD11bhigh F4/80intermediate.

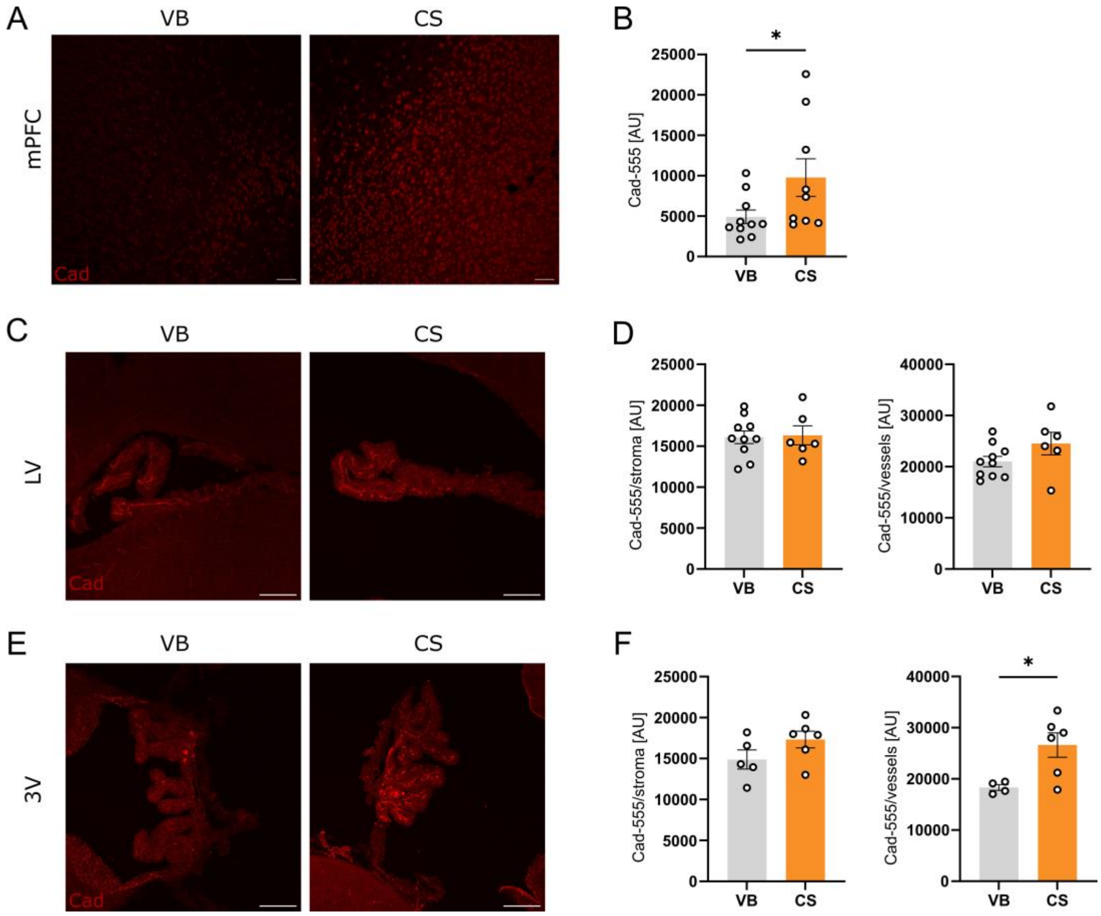


Fig. S5. Mode of delivery affects brain barrier functional permeability. (A) Representative images of tracer extravasation in the mPFC of P8 pups vaginally born (VB) and delivered by cesarean section (CS) after an *in vivo* Cadaverine Alexa Fluor 555 (1kDa) i.p. injection, scale bar = 50µm. (B) Quantification of fluorescence intensity in the mPFC (mean ± SEM; Mann-Whitney, *$p<0.05$, VB n=10, CS n=9). (C) Representative images of tracer in the lateral ventricle (LV), scale bar = 100µm. (D) Quantification of fluorescence intensity in the stroma and the vessels of the LV (mean ± SEM; t-test, VB n=10, CS n=6). (E) Representative images of tracer in the third ventricle (3V), scale bar = 100 µm. (F) Quantification of fluorescence intensity in the stroma and the vessels of the 3V (mean ± SEM; t-test, *$p<0.05$, VB n=4-5, CS n=6).

**Table S1.** Key antibody references.

| ANTIBODY | SOURCE | IDENTIFIER | APPLICATION | DILUTION |
|---|---|---|---|---|
| **Myeloid cells** | | | | |
| PerCP-conjugated anti-CD11b | Biolegends | Cat#101230 | Flow cytometry | 1:50 |
| Brilliant Violet 785-conjugated anti-CD45 | Biolegends | Cat#103149 | Flow cytometry | 1:50 |
| APC-conjugated anti-LY-6C | Miltenyi | Cat#130-123-796 | Flow cytometry | 1:100 |
| BV421-conjugated anti-CX3CR1 | Biolegends | Cat#149023 | Flow cytometry | 1:50 |
| PE-Cyanine7-conjugated anti-F4/80 | Biolegends | Cat#123114 | Flow cytometry | 1:50 |
| Brilliant Violet 510-conjugated anti-IA-IE | Biolegends | Cat#107636 | Flow cytometry | 1:50 |
| Rabbit anti-CD68 | Invitrogen | Cat#PA5-78996 | Immunofluorescence | 1:150 |
| Guinea pig anti-IBA1 | Synaptic Systems | Cat#234-308 | Immunofluorescence (in the choroid plexus) | 1:500 |
| Rabbit anti-IBA1 | Fujifilm | Cat#019-19741 | Immunofluorescence (in cortex) | 1:500 |
| **T cells** | | | | |
| Brilliant Violet 605-conjugated anti-CD4 | Biolegends | Cat#100548 | Flow cytometry | 1:50 |
| Brilliant Violet V785-conjugated anti-CD45 | Biolegends | Cat#103149 | Flow cytometry | 1:50 |
| PerCP-conjugated anti-CD11b | Biolegends | Cat#101230 | Flow cytometry | 1:50 |
| **Tight junctions** | | | | |
| rabbit anti-ZO-1 | Invitrogen | Cat#61-7300 | Immunofluorescence | 1:150 |
| **Vessels** | | | | |
| Rat anti-PV1 | BD Biosciences | Cat#550563 | Immunofluorescence | 1:50 |
| Goat anti-podocalyxin | R&D | Cat#AF1556 | Immunofluorescence | 1:150 |
| Rabbit anti-Aquaporin 4 | Millipore | Cat#AB2218 | Immunofluorescence | 1:200 |
| Isolectine GS-IB4, Alexa Fluor™ 647 conjugated | Invitrogen | Cat#I32450 | Immunofluorescence | 1:200 |
| **Secondary antibodies** | | | | |
| Donkey anti-goat Alexa Fluor™ Plus 405 | Invitrogen | Cat#A48259 | Immunofluorescence | 1:200 |
| Donley anti-goat Alexa Fluor™ Plus 647 | Invitrogen | Cat#A32849 | Immunofluorescence | 1:200 |
| Donkey anti-rat Alexa Fluor™ Plus 647 | Invitrogen | Cat#A48272 | Immunofluorescence | 1:200 |

| | | | | |
|---|---|---|---|---|
| Donley anti-guinea-pig Alexa Fluor™ Plus 647 | Jackon Immunores earch | Cat#706-605-148 | Immunofluorescence | 1:200 |
| Donkey anti-rabbit Alexa Fluor™ Plus 488 | Invitrogen | Cat#A32790 | Immunofluorescence | 1:200 |

### 3. REFERENCES

1. Champagne, F.A., Curley, J.P., Keverne, E.B., and Bateson, P.P.G. (2007). Natural variations in postpartum maternal care in inbred and outbred mice. Physiol. Behav. *91*, 325–334. https://doi.org/10.1016/j.physbeh.2007.03.014.
2. Dominguez-Belloso, A., Schmutz, S., Novault, S., Travier, L., and Deczkowska, A. (2022). Isolation and characterization of the immune cells from micro-dissected mouse choroid plexuses. J. Vis. Exp. https://doi.org/10.3791/63487.
3. Schindelin, J., Arganda-Carreras, I., Frise, E., Kaynig, V., Longair, M., Pietzsch, T., Preibisch, S., Rueden, C., Saalfeld, S., Schmid, B., et al. (2012). Fiji: an open-source platform for biological-image analysis. Nat. Methods *9*, 676–682. https://doi.org/10.1038/nmeth.2019.
